# A shared RNA structural element mediates hnRNP K binding by two divergent lncRNAs in the DNA damage response

**DOI:** 10.1101/2025.07.24.666508

**Authors:** Salvatore Assenza, Constanza Blanco, Cyril Favard, Julie Carnesecchi, Marco Marcia, Isabel Chillón

## Abstract

LncRNAs are typically poorly conserved at the sequence level, which complicates the use of sequence-based approaches to identify conserved functions. The stress-associated lincRNA-p21 is a paradigm for a sequence-divergent lncRNA. In mice, the interaction between lincRNA-p21 and the hnRNP K protein activates the *Cdkn1a* gene *in cis*, encoding the cell-cycle regulator p21, a key effector of the cellular stress response to DNA damage. However, the molecular details of this interaction, and its conservation in other species, remained unknown. To address this, we applied an integrated structural and functional approach, combining *in silico*, *in vitro*, *ex cellulo*, and *in cellulo* assays. We identified a structural element in mouse and human lincRNA-p21 containing a tetranucleotide motif conserved across more than 100 mammals. This element directly mediates a weak binding to the hnRNP K protein in both species, which may enable rapid and reversible modulation of *Cdkn1a* and timely resolution of the DNA damage response. Conserved hnRNP K binding in humans suggests that the mechanism of *Cdkn1a* activation may also occur in this species. Our findings establish a shared RNA-protein contact between two syntenic but sequence-divergent transcripts, offering a general strategy for identifying functional RNA elements in other lncRNAs with limited sequence conservation.

## INTRODUCTION

Eukaryotic long non-coding RNAs (lncRNAs) are critical regulators of transcription and gene expression ^1^, yet have long been dismissed as junk due to pervasive transcription and poor linear sequence conservation ^2^. Where function has been assessed, this lack of conservation has led to species-specific characterizations, obscuring evolutionary significance. ^3^. RNA structure has emerged as a primary driver of lncRNA function ^4^. Examples include the MEG3 pseudoknot, which activates the p53 pathway, and the MALAT1 triple helix, which confers stability against degradation ^5,6^. However, given the complexity of identifying such motifs across species when sequence conservation is low, a single technique is rarely sufficient and often requires complementary structural approaches.

LincRNA-p21 (Trp53cor1) shows low global sequence conservation in mammals. Its mouse and human orthologs are syntenic but diverge in sequence, gene structure, and function. Mouse lincRNA-p21 (mlincRNA-p21) is a 3-kb spliced, polyadenylated transcript (RefSeq: NR_036469), transcribed antisense to and upstream of *Cdkn1a*, encoding the DNA damage response effector p21 (Fig. 1A) ^7^. The human ortholog (hlincRNA-p21) lies in a syntenic region and produces two unspliced transcripts of 3 kb (GenBank: KU881769) and 4 kb (GenBank: KU881768), sharing a common 5’ end (Fig. 1A-B) ^8^. Both are canonical p53 targets. MlincRNA-p21 is strictly nuclear and regulates *Cdkn1a in cis* by binding hnRNP K, an ssDNA/RNA-binding protein that promotes p53-dependent *Cdkn1a* transcription activation ^9,10^. HlincRNA-p21, in contrast, localizes to both the nucleus and the cytoplasm. Nuclear hlincRNA-p21 also binds hnRNP K, though its functional relevance remains unexplored ^11^.

**Figure 1.**
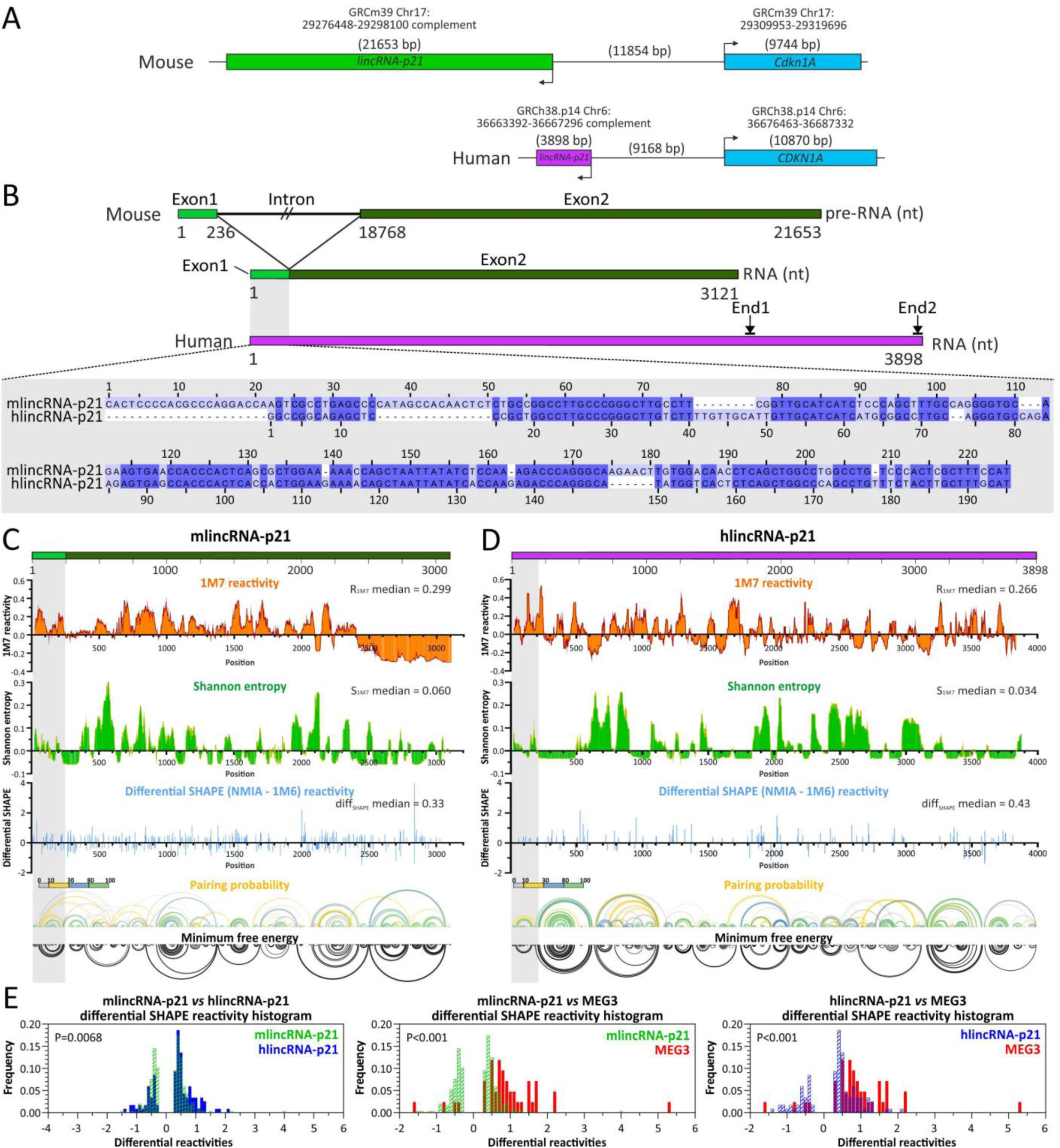
The structural architecture of mlincRNA-p21 and hlincRNA-p21 reveals a decentralized RNA organization. **A.** Schematic representation of the syntenic mouse and human *lincRNA-p21* genes. Gene lengths and distances are shown at an approximate scale. **B.** Schematic representation of the mouse and human lincRNA-p21 transcripts. A grey box indicates the region displaying high sequence similarity between mlincRNA-p21 and hlincRNA-p21. A grey inset displays the sequence alignment of mlincRNA-p21 and hlincRNA-p21 generated using MUSCLE. **C** and **D.** Structural architecture of mlincRNA-p21 by SHAPE-MaP (C) and hlincRNA-p21 by SHAPE (D). 1M7 reactivity (top panel) and Shannon entropy (second panel from the top) values are shown as the median reactivity over 55-nt sliding windows relative to the global median. Differential SHAPE values (NMIA – 1M6) as obtained using the previously described differential SHAPE scripts ^15^ (third panel from the top). Base-pairing probabilities are indicated by arcs connecting base pairs and are color-coded based on probability (fourth panel from the top). Minimum free energy secondary structure model (bottom panel). **E.** Histogram showing the distribution of differential SHAPE reactivity values. P-value obtained by distribution comparison using a Kruskal-Wallis test.

Understanding the molecular basis of the lincRNA-p21/hnRNP K interaction is essential given p21’s central role in the DNA damage response. This raises a key question: how can two syntenic but divergent lncRNAs, differing in sequence and localization bind orthologous proteins? To address this, we combined *in vitro*, *ex cellulo*, and *in cellulo* chemical probing to identify a structural element localized at the 5’-end of both mlincRNA-p21 and hlincRNA-p21, which contained a sequence motif conserved across more than 100 species. *In silico* analysis, molecular dynamics simulations, and *in vitro* binding assays showed that this structural element mediates binding to the KH3 domain of hnRNP K, a finding further supported by functional assays in cells. Conservation of this interaction between mouse and human suggests a shared mechanism for *Cdkn1a* activation, which has been so far demonstrated only in mice. Our work provides a molecular basis for an lncRNA-protein interaction that fine-tunes *Cdkn1a* expression during the cellular stress response to DNA damage and offers a general strategy for discovering functional RNA motifs in poorly conserved lncRNAs.

## RESULTS

### Mouse and human lincRNA-p21 display a dispersed, highly structured architecture

To search for structural RNA motifs in mlincRNA-p21 and hlincRNA-p21, we generated structural maps by Selective 2′-Hydroxyl Acylation analyzed by Primer Extension (SHAPE) and by SHAPE analyzed by mutational profiling (SHAPE-MaP) (Fig. 1C-D). Both techniques rely on chemical probes that acylate the 2′-OH of structurally flexible, typically single-stranded, residues, providing single-nucleotide resolution ^12^. We produced spliced mlincRNA-p21 and the hlincRNA-p21 long isoform (Fig. 1B) by T7 *in vitro* transcription under non-denaturing conditions, which preserve co-transcriptional folding ^13^. This is important given lincRNA-p21’s 3-4 kb length and its potential to adopt multiple secondary/tertiary configurations, as we previously observed for HOTAIR ^14^.

RNA was chemically probed with 1M7, 1M6, and NMIA. Differential SHAPE (1M6 minus NMIA) captured variations in the flexibility of backbone residues arising from noncanonical and tertiary interactions. This was incorporated, together with 1M7 reactivity, into partition function calculations to derive base-pairing probabilities and Shannon entropy (Fig. 1C-D) ^15–17^. All three reagents showed high reproducibility (Fig. S1A-B), and treated samples showed significantly elevated raw signal over DMSO controls, confirming robust structural signal (Fig. S2A-B).

Both transcripts showed a globally low median 1M7 reactivity (<0.4), comparable to other structured lncRNAs (MEG3, XIST) ^5,18^ and viral genomes (HCV, HIV) ^19,20^, indicating extensive base pairing (Fig. 1C-D, upper panel). The global median Shannon entropy was similarly low, comparable to that of HCV ^19^, indicating a high probability of folding into a single dominant conformation (Fig. 1C-D, second panel). Differential SHAPE analysis revealed few nucleotides with low but statistically significant differential reactivity in hlincRNA-p21 and mlincRNA-p21 (Fig. 1C-D, third panel), although the two distributions differed significantly from each other (Fig. 1E). Still, this difference was minor relative to MEG3, which is one of the few lncRNAs known to adopt a compact 3D fold, characterized by a small number of strong differential SHAPE signals (Fig. 1E). By contrast, the lincRNA-p21 profiles suggest neither ortholog forms stable tertiary nor noncanonical interactions. Consistent with this, pairing probability and minimum free energy diagrams for both transcripts showed short, highly structured motifs (low SHAPE reactivity, low Shannon entropy) distributed across the full length of the transcript, without higher-order hierarchical organization (Fig. 1C-D, fourth panel).

In summary, our SHAPE and SHAPE-MaP data support a decentralized, dispersed organization of lincRNA-p21, comprising discrete, highly structured RNA motifs distributed across the transcript.

### The structure of the lincRNA-p21 5’-end region comprises a putative hnRNP K binding motif

An RNA pull-down analysis with truncated mlincRNA-p21 constructs previously localized hnRNP K binding to the first 780 nt ^7^. We reasoned that the binding motif should lie within a region of shared sequence or structure between mouse and human transcripts. We identified high sequence similarity in the first 225 nt of mlincRNA-p21 (exon 1) and the first 195 nt of hlincRNA-p21 (Fig. 1B, grey inset).

Using our SHAPE data, we modeled the secondary structure of both transcripts with the Superfold pipeline and examined their 5’ regions (Fig. 2A-B) ^16,21^. The SHAPE reactivity was consistent with the resulting models (Fig. S2C-D). Mouse exon 1 folds as a single independent structural unit (helices H1-H10) and ends in a four-way junction (Fig. 2A). The corresponding human region does not form one continuous unit. It contains two hairpins, a short segment (helices H2-H3) ending in a four-way junction, and the 5’ strand of an independent motif (pos. 172-195) (Fig. 2B). The four-way junction itself, however, has a similar architecture in both species, each with three short helical arms. Notably, most nucleotides with differential 1M6/NMIA reactivity, indicating base stacking or local tertiary interactions, respectively, map to this four-way junction. This suggests that the four-way junction adopts a complex 3D configuration, potentially orienting a protein-binding motif.

**Figure 2.**
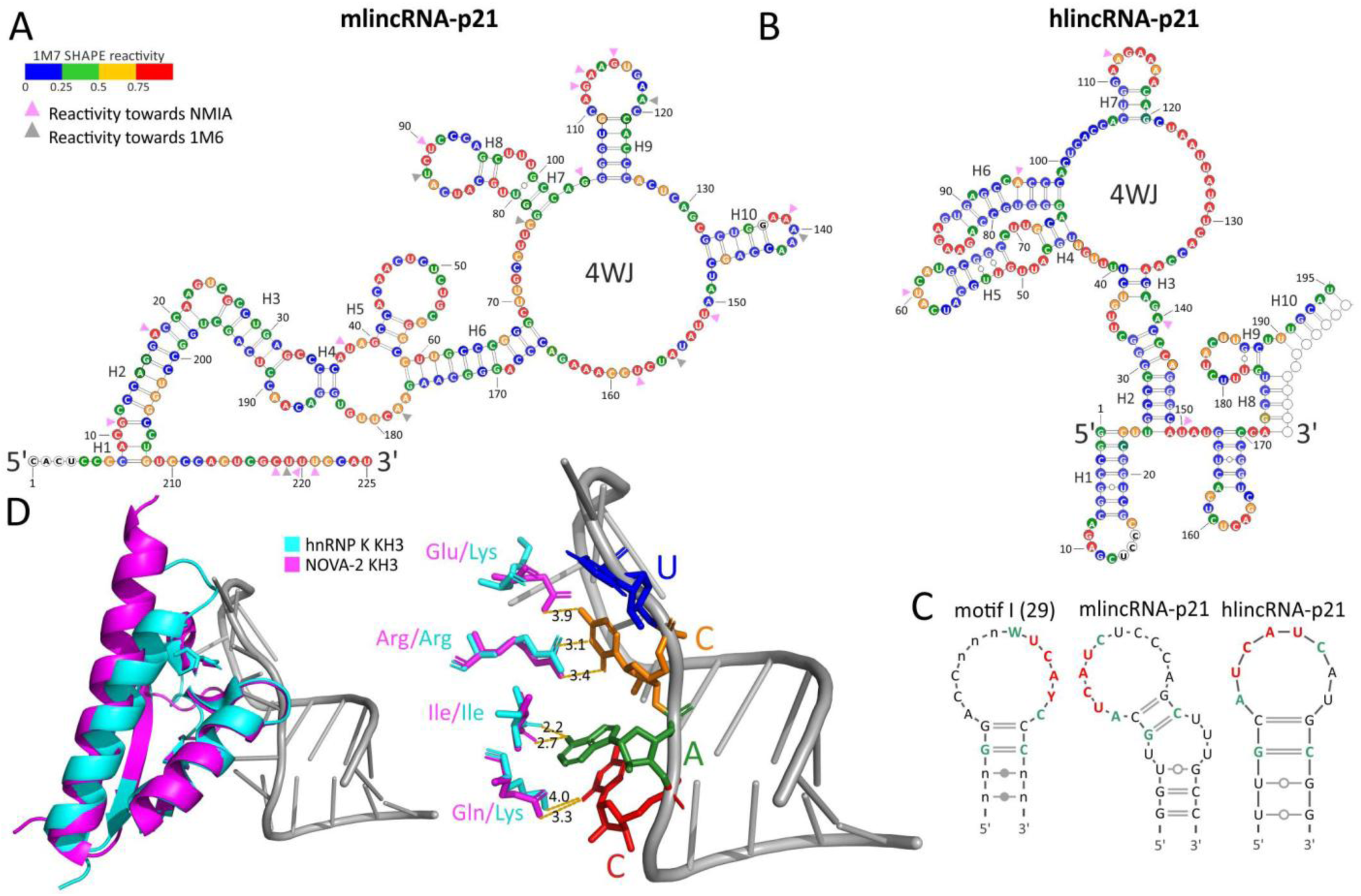
A putative hnRNP K binding RNA structure is present in mlincRNA-p21 and hlincRNA-p21. A, **B.** Secondary structure models of mlincRNA-p21 positions 1-225 **(A)** and hlincRNA-p21 positions 1-195 **(B)** obtained with the software Superfold, using the SHAPE-MaP and SHAPE reactivity values as constraints for minimum free energy calculations. **C.** Schematic representation of the putative hnRNP K binding RNA hairpin as previously described by SELEX (left panel), and the ones found in mlincRNA-p21 (middle panel) and hlincRNA-p21 (right panel). Essential protein recognition nucleotides are depicted in red; conserved nucleotides fine-tuning protein affinity are depicted in green. **D.** Superposition of the crystal structures of the NOVA-2 KH3 domain bound to the 20-mer RNA hairpin (PDB 1EC6) and the hnRNP K KH3 domain in complex with a 6-mer ssDNA (not displayed) (PDB 1ZZI). Hydrogen bonds established between the conserved UCAC tetranucleotide and each of the protein KH3 domains are indicated in Angstroms (Å) (right panel).

Within this junction, the first helical arm of each transcript (mouse: 78-102; human: 45-74) closely resembles a SELEX-derived RNA element that binds the KH3 domains of NOVA-1/NOVA-2 ^22,23^ (Fig. 2C). This element comprises a UCAY tetranucleotide in the loop of a 20-nt hairpin, flanked by a conserved W and C, plus an ACCNNN stretch that contributes to affinity. Both lincRNA-p21 orthologs contain the essential UCAY tetranucleotide and the flanking W/C positions (Fig. 2C) but lack the ACCNNN stretch, and their first Watson-Crick pair (C-G) differs from the SELEX consensus (G-C).

To assess whether hnRNP K’s KH3 domain could potentially bind this hairpin, we compared KH domain sequences and confirmed that hnRNP K KH3 is more similar to the NOVA-1/2 KH3 domain than to its own KH1/KH2 domains (Fig. S3A-B) ^24^. Superimposing the NOVA-2 KH3–RNA hairpin structure (1EC6) ^25^ with the hnRNP K KH3–ssDNA structure (1ZZI) ^26^ gave excellent agreement (RMSD = 0.911 Å), with only the C-terminal α’ helix and variable loop shorter in hnRNP K. Five of eight RNA-contacting residues are conserved between NOVA-2 and hnRNP K (Fig. S3A). Of the four substituted contacts, most preserve the interaction type: at the first UCAY position, van der Waals and π-π stacking contacts are unaffected by Gly18/Ala19 to Ser27; at the second position, the cytidine loses one hydrogen bond (Glu14 to Lys22), weakening this contact; at the third position, adenine’s hydrogen bond with the Ile-41 backbone is maintained, while van der Waals contacts with Leu-21/Leu-28 are preserved *via* the hydrophobic substitutions Ile-29/Ile-36; at the fourth position, the pyrimidine’s hydrogen bond with Gln-40 (NOVA-2) is replaced by one with Lys-48 (hnRNP K), and although lincRNA-p21 has a uridine rather than cytidine here, the shared pyrimidine oxygen preserves this contact (Fig. 2D, S3A).

Together, these structural and sequence comparisons suggest a conserved UCAY motif within a terminal stem-loop, potentially recognized by hnRNP K’s KH3 domain, albeit with a weaker interaction than in the NOVA paradigm.

### The 5’-end region of lincRNA-p21 is conserved across mammals

We next asked whether the putative hnRNP K-binding helical arm, located within the four-way junction of the mlincRNA-p21 and hlincRNA-p21 5’ region, is phylogenetically conserved. Using a locally built BLASTN database, we searched 111 mammalian genomes for homology with mlincRNA-p21 (pos. 1-225) and hlincRNA-p21 (pos. 1-195). The 5’-end of lincRNA-p21 is present in nearly all mammals except monotremes and marsupials, with high sequence similarity despite group-or species-specific insertion gaps (Fig. S4). The KH3 recognition motif UCAY is widely conserved, typically as UCAU, but is absent in guinea pigs and mutated to UCAG in bats, suggesting these species lack hnRNP K binding *via* this motif or have no hnRNP K binding.

A higher primate-specific insertion (Fig. S4, yellow) falls within the putative hnRNP K-binding helical arm and is a critical structural part of it (Fig. 3A, C, yellow). This insertion lengthens the helical arm in higher primates and shifts the base pairing (H4-5 in humans vs. H7-8 in mice) (Fig. 3B-C), suggesting that the human helical arm is an evolutionarily recent variant. In contrast, the mouse structure may represent the ancestral form compatible with most mammalian sequences, including primates.

**Figure 3.**
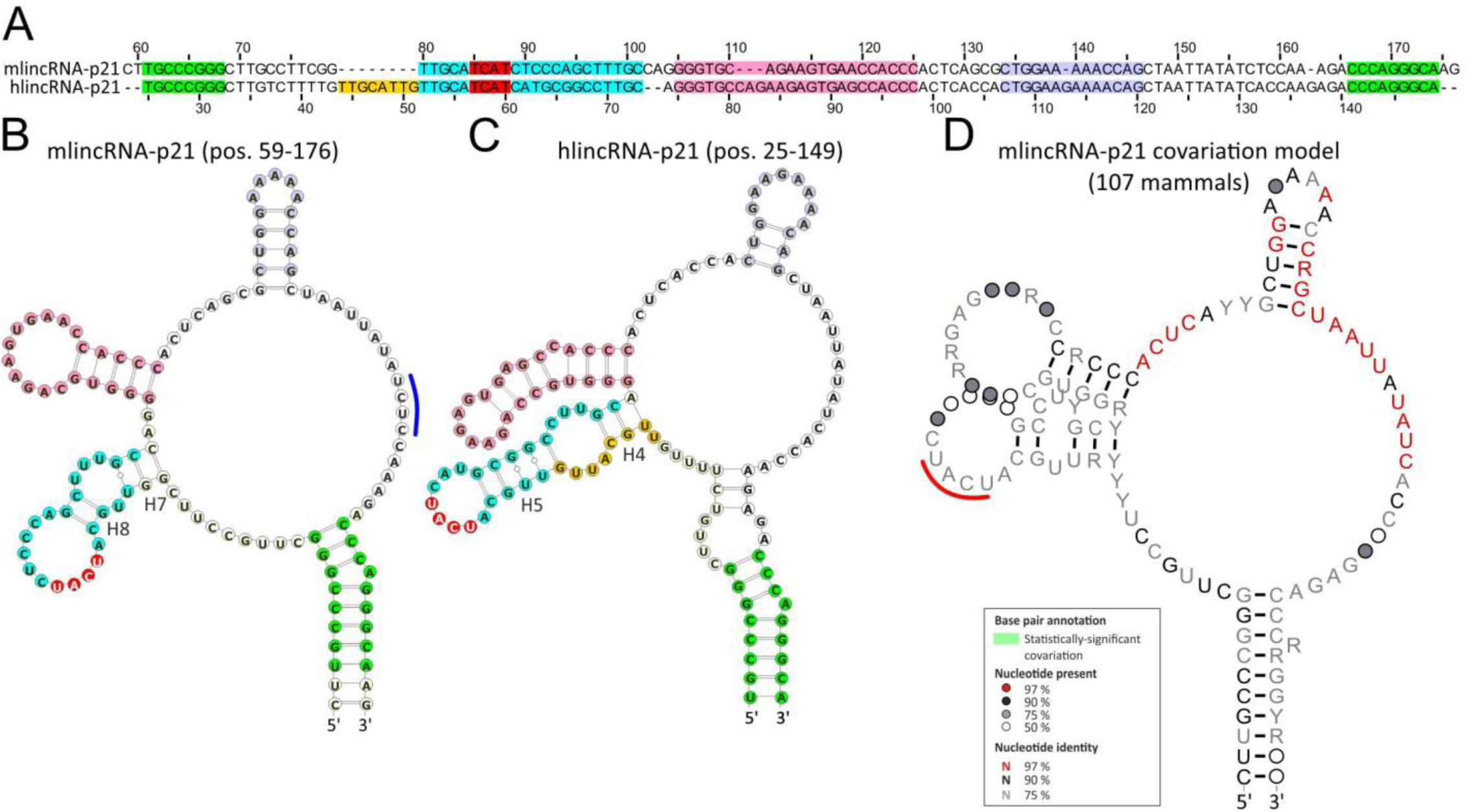
The lincRNA-p21 four-way junction is highly conserved in sequence among mammals. **A.** Detail of the sequence alignment between mlincRNA-p21 and hlincRNA-p21 (Fig. S4). Regions corresponding to the different structural motifs are highlighted in colors. The UCAY tetranucleotide is indicated in red. The higher-primate-specific sequence insertion is colored in yellow. **B** and **C**. Schematic representation of the mlincRNA-p21 (B) and hlincRNA-p21 (C) four-way junction showing structural motifs color-coded as in A. A blue line indicates a putative competing sequence with the UCAU located in the H7-H8 helical arm of mlincRNA-p21. **D.** Schematic representation of the consensus four-way junction structure obtained by structural alignment of 107 mammalian lincRNA-p21 homologous sequences using the Infernal software. A red line indicates the position of the UCAY tetranucleotide. A covariation model obtained by a seed-alignment of 42 species to the mlincRNA-p21 structure was used. The statistical significance of the covariation analysis was assessed by R-scape for each helical arm (P=0.1).

To test this, we built a covariation model from a 42-species seed alignment and the mlincRNA-p21 four-way structure using Infernal. We searched 111 mammal species and successfully aligned 107 to our mouse covariation model (Fig. 3D). This confirmed strong sequence conservation, with up to 97% nucleotide identity in the single-stranded region flanking the H10 hairpin and in the hairpin itself. The UCAY motif was conserved as UCAU in 75% of species. However, R-scape found no statistically significant structural covariation, likely due to insufficient substitution events given the high sequence conservation, limiting analytical power. However, we cannot rule out the possibility that the putative hnRNP K-binding helical arm structure is present in mammals beyond mice and humans. Indeed, despite the higher-primate insertion, a minimal structural element comprising the terminal stem-loop of the helical arm, flanked by two consecutive G-C pairs, as identified by SELEX, is conserved between mouse and human and may be conserved across all mammals that retain the UCAY motif.

In conclusion, sequence and structural conservation analysis of the putative hnRNP K-binding helical arm across more than 100 mammalian species revealed marked conservation of a UCAU tetranucleotide in a terminal stem-loop and its flanking helix, suggesting the structure itself may also be conserved, despite the lack of statistically significant covariation.

### *In silico* models suggest stable interaction between lincRNA-p21 and hnRNP K

To inspect the stability of our experimentally determined UCAU-containing helical arms, we performed microsecond-long atomistic molecular dynamics simulations. Initial 3D models were generated with simRNA ^27,28^ from our mlincRNA-p21 and hlincRNA-p21 secondary structures, then simulated using AMBER ^29^. Both free structures remained overall stable (Fig. 4A-B, yellow bars). Notably, a novel A-48:U-71 pair emerged at the internal loop of the hlincRNA-p21 helical arm.

**Figure 4.**
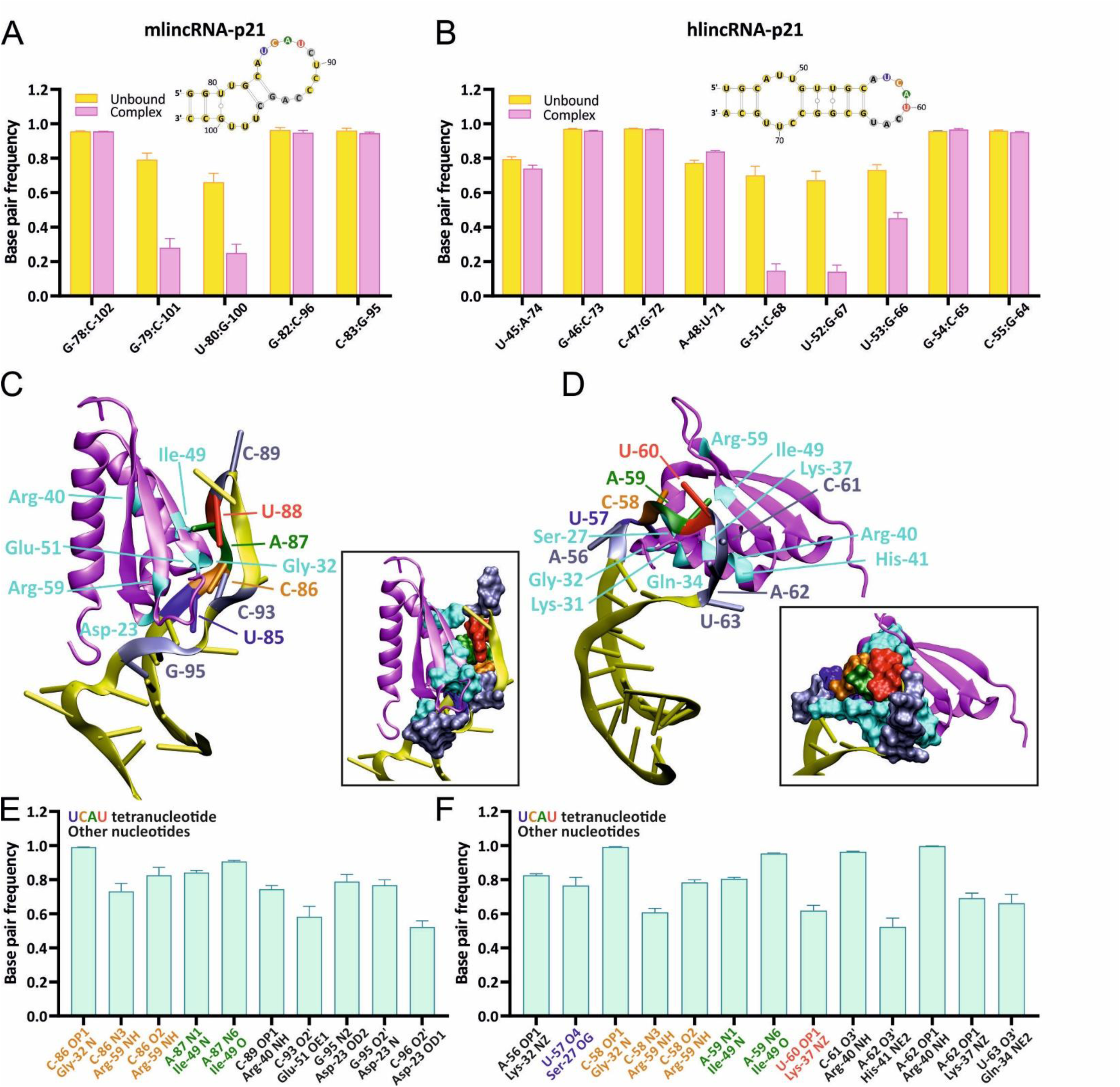
LincRNA-p21 forms an extended network of hydrogen bonds with the KH3 domain. A, **B.** Census of the base pairs detected in the RNA molecules for mlincRNA-p21 **(A)** and hlincRNA-p21 **(B)**. The labels in the horizontal axis indicate the base pairs under consideration; in the case of hlincRNA-p21, the underlined label indicates the A-48:U-71 base pair observed in the simulations but not reported by the SHAPE experiments. The yellow and magenta bars correspond to the populations observed for free RNA and the RNA bound to the hnRNP K KH3 domain, respectively. The secondary structures of the RNA fragments are represented in the insets, color-coded as as in panels C and D (see below). **C, D.** Representative snapshots of the complexes formed by the hnRNP K KH3 domain and mlincRNA-p21 **(C)** or hlincRNA-p21 **(D)**. The overall structure of the two molecules is depicted as cartoons, with magenta and yellow used to represent the protein and the RNA, respectively. Residues directly participating in the formation of hydrogen bonds are highlighted by the use of a different color: the UCAU tetramer is colored in blue, orange, green and red following the 5’ to 3’ direction; nucleosides outside the UCAU tetramer that participate in hydrogen bonds are represented in light grey; amino acids partners for the hydrogen bonds are drawn in cyan. All the highlighted residues are annotated with a label of the same color. Insets in C and D represent the same snapshots, where the hydrogen-forming residues are depicted using a space-filling representation to convey their spatial occupancy better. **E, F.** Population analysis of the hydrogen bonds formed between lincRNA-p21 and the hnRNP K KH3 domain for mlincRNA-p21 **(E)** and hlincRNA-p21 **(F)**. Only the most frequent (>50%) hydrogen bonds are reported in the plot. The labels on the horizontal axis indicate the residues and atoms participating in the hydrogen bond, with the tetramer residues highlighted with the same color code as in panels C and D.

We next modeled the hnRNP K KH3–RNA interaction, starting from the KH3 crystal structure (PDB 1ZZI) ^26^, with the ssDNA dTCCC replaced by rUCAU, then aligned to pre-equilibrated mouse and human helical arms. Binding to KH3 reduced Watson-Crick pairing frequency overall (2.8-fold in mouse, 4.8-fold in human; magenta bars, Fig. 4A-B), though most individual base pairs were only marginally affected (1.1- to 1.6-fold), indicating overall structural stability within the complex. A few pairs were strongly perturbed instead, such as G79:C101 and U80:G100 in mouse (3- to 4-fold), and G51:C68, U52:G67, and U53:G66 in human (up to >6-fold), while the novel A48:U71 pair showed a mild increase. Perturbed pairs were consistently distant from the binding interface, unlike closer, unaffected pairs.

Simulations revealed RNA-protein contacts extending beyond the UCAU tetranucleotide itself (Fig. 4C-D), involving both hydrogen bonds (Fig. 4E-F) and van der Waals interactions (Fig. S5A-B), located within the SELEX-defined minimal stem-loop. The core shared sequence motif comprises five hydrogen bonds from the second (C) and third (A) UCAU residues to Gly-32, Ile-49, and Arg-59. Two of these, C:Arg-59 and A:Ile-49, matched our crystal structure superimposition, while an unanticipated backbone-backbone C:Gly-32 contact was the most frequent overall. The predicted U:Lys-48 hydrogen bond was rare (5% in mouse, 12% in human), though our van der Waals analysis showed that these residues remain in nonspecific proximity (∼13 and 21 contacts, respectively; Fig. S5A-B).

Despite this shared core, sequence and structural differences elsewhere in the helical arm led to divergent binding, where the human helical arm engaged more residues overall (8 nucleosides, 9 amino acids) than the mouse (6 nucleosides, 6 amino acids) and formed more hydrogen bonds on average (10 vs. 7.6; Fig. S5C), suggesting a more stable hnRNP K KH3 interaction for hlincRNA-p21.

In summary, molecular dynamics simulations aligned with our experimental secondary structures and revealed a rich, partly novel network of RNA-protein contacts beyond the core UCAY motif, with nearby nucleosides modulating binding in a system-dependent manner, and hlincRNA-p21 appearing to bind more stably than mlincRNA-p21.

### HnRNP K binds lincRNA-p21 *in vitro* through its KH3 RNA-binding domain

To determine whether hnRNP K’s KH1, KH2, or KH3 domain mediates lincRNA-p21 binding, we performed an *in vitro* protein-RNA interaction assay with UV crosslinking and RNase A digestion (Fig. 5, S6). MBP-tagged full-length (FL) hnRNP K or each isolated KH domain was incubated with fluorescently labeled short mlincRNA-p21 (first 211 nt, WT-short-mlincRNA-p21) and hlincRNA-p21 (first 152 nt, WT-short-hlincRNA-p21) transcripts (Fig. 5A-B). After crosslinking and RNase digestion, protected RNA was visualized on denaturing gels (Fig. 5C-D, top) alongside protein content, as determined by Coomassie staining (Fig. 5C-D, second panel). Both transcripts bound FL hnRNP K, in agreement with previous observations ^7,11^. Neither transcript bound KH1 nor KH2, but both bound KH3 (Fig. 5C-D, blue bars), confirming our hypothesis. KH3 binding alone did not fully recapitulate FL binding, suggesting that neighboring protein sequences confer additional affinity, as previously shown for NOVA-2 ^22^.

**Figure 5.**
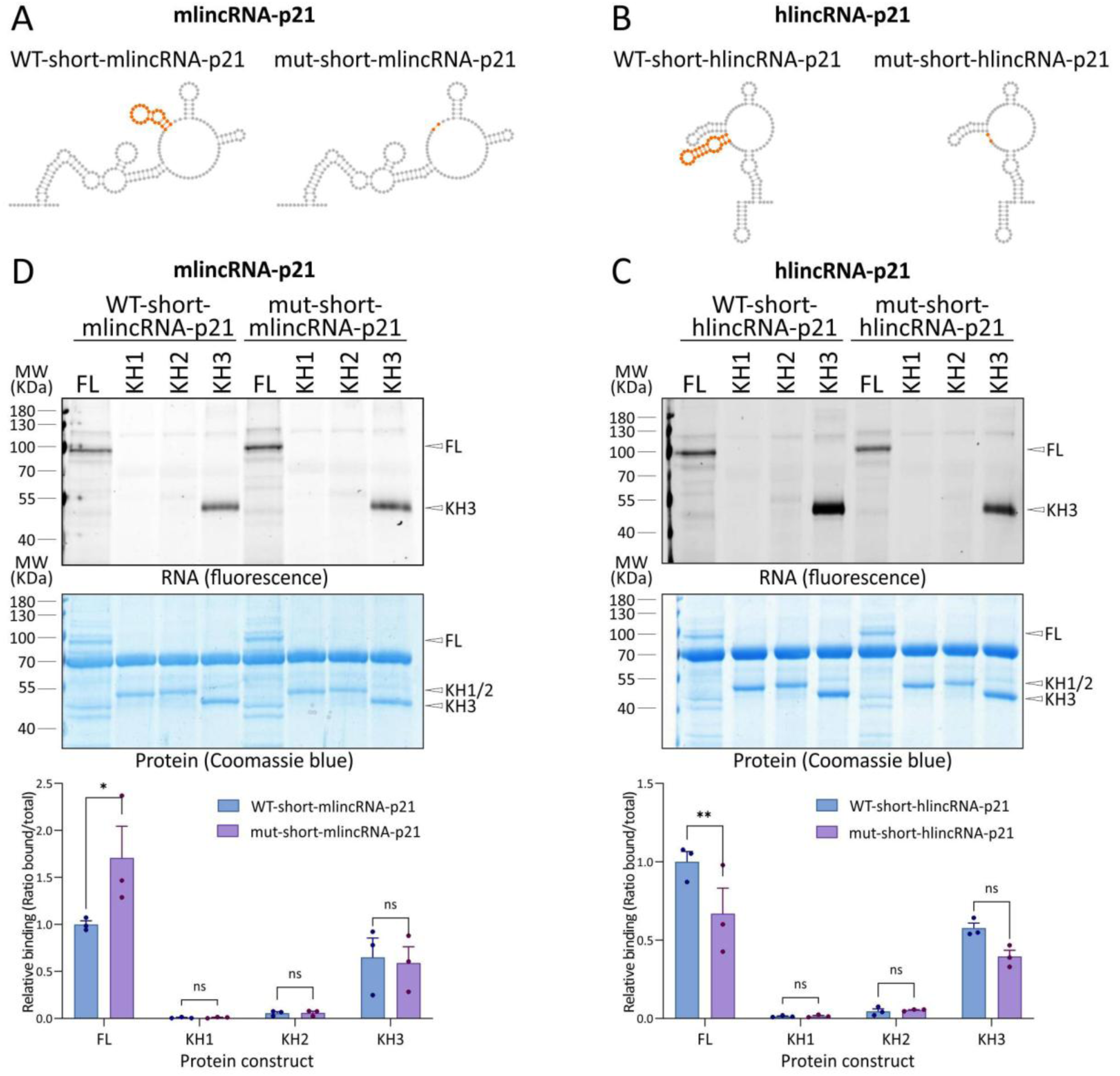
The KH3 domain of hnRNP K binds lincRNA-p21. A and. **B.** Schematic representation of the 5’-end wild-type structures (left) of mlincRNA-21 (A) and hlincRNA-21 (B) and the resulting deletion of the UCAY-containing helical arm (right, highlighted in orange in the wild-type construct). **C and D**. Fluorescent protein-RNA interaction assay followed by UV-crosslinking and RNase digestion, performed *in vitro* with purified pETM44-hnRNPK-FL protein or the individual KH domains (pET-MBP-hnRNPK-KH1, pET-MBP-hnRNPK-KH2, and pET-MBP-hnRNPK-KH3), and short hlincRNA-p21 (C) and mlincRNA-p21 (D) constructs. Interactions were detected on denaturing SDS-PAGE gels by Cy5-UTP signal (upper panels). Coomassie-stained gels reveal the protein content (middle panels). BSA is visible at 70 kDa on the Coomassie gel, and a ladder is presented to underscore the corresponding molecular weight. Quantification of relative RNA-binding of the isolated KH domains was compared to the FL hnRNP K protein (lower panels). The graph represents three independent assays. Statistical test by two-way ANOVA (*: P=0.05, **: P=0.01).

To test the functional significance of the UCAU-containing helical arm, we generated mutant short constructs lacking this element for both species (mut-short-mlincRNA-p21, mut-short-hlincRNA-p21; Fig. 5A-B). Human WT and mutant constructs showed significantly different binding to FL hnRNP K (Fig. 5C). However, a similar trend with KH3 alone did not reach significance, likely reflecting KH3’s generally weaker binding relative to the FL protein. Counterintuitively, the mouse mutant bound hnRNP K more strongly than WT (Fig. 5D), potentially because the removal of the helical arm exposes a downstream competing sequence (UCUC, positions 156-159) with higher intrinsic affinity.

In summary, hnRNP K binds both mlincRNA-p21 and hlincRNA-p21 via its KH3 domain, not KH1 or KH2. The UCAU-containing helical arm mediates hlincRNA-p21 binding *in vitro*. For mlincRNA-p21, we could not confirm this *via* the same element, but we cannot rule out a competing higher-affinity sequence that may or may not be accessible in cells.

### The lincRNA-p21 UCAU-containing helical arms are protected by proteins inside cells

To determine whether the 5’-end regions of mlincRNA-p21 and hlincRNA-21 are involved in protein binding inside cells, we performed a ΔSHAPE assay, which relies on the nucleotide flexibility changes that occur on the RNA at RNA-protein binding sites ^30^. ΔSHAPE was obtained by subtracting *in cellulo* SHAPE reactivity values (inside cells in the presence of proteins) from *ex cellulo* reactivity values (RNA extracted and deproteinized before NAI chemical probing). Because endogenous transcript levels were too low for efficient SHAPE-MaP amplification ^16^, we transfected HCT116 (human) and NIH-3T3 (mouse) cells with hlincRNA-p21 or mlincRNA-p21 expression plasmids, respectively.

*Ex cellulo* reactivity correlated well across replicates for both transcripts (Fig. S7A-B, left), but *in cellulo* replicates showed poor correlation (Fig. S7A-B, right), despite all replicates passing quality control (Fig. S7C-D). This indicates that deproteinized lincRNA-p21 adopts a single, reproducible structure, whereas each *in cellulo* replicate captures a distinct protein-occupancy state. Given this variability, we calculated ΔSHAPE separately for each *in cellulo* replicate against average *ex cellulo* values (Fig. S8), then averaged these to identify statistically significant, reproducible changes (Fig. 6A-B). This identified three consistent regions in mlincRNA-p21 and five in hlincRNA-p21 with the largest significant ΔSHAPE changes (Fig. 6A-B, grey/dotted boxes). These regions comprise residues with positive ΔSHAPE values, indicating protection from modification in the cellular environment, and negative ΔSHAPE values, denoting enhanced reactivity in cells due to conformational changes upon protein binding. In both cases, the ΔSHAPE analyses detected the UCAU-containing helical arms in both mlincRNA-p21 at positions 75-104 (shaded in grey in Fig. 6A and C) and hlincRNA-p21 at positions 50-58 (shaded in grey in Fig. 6B and D).

**Figure 6.**
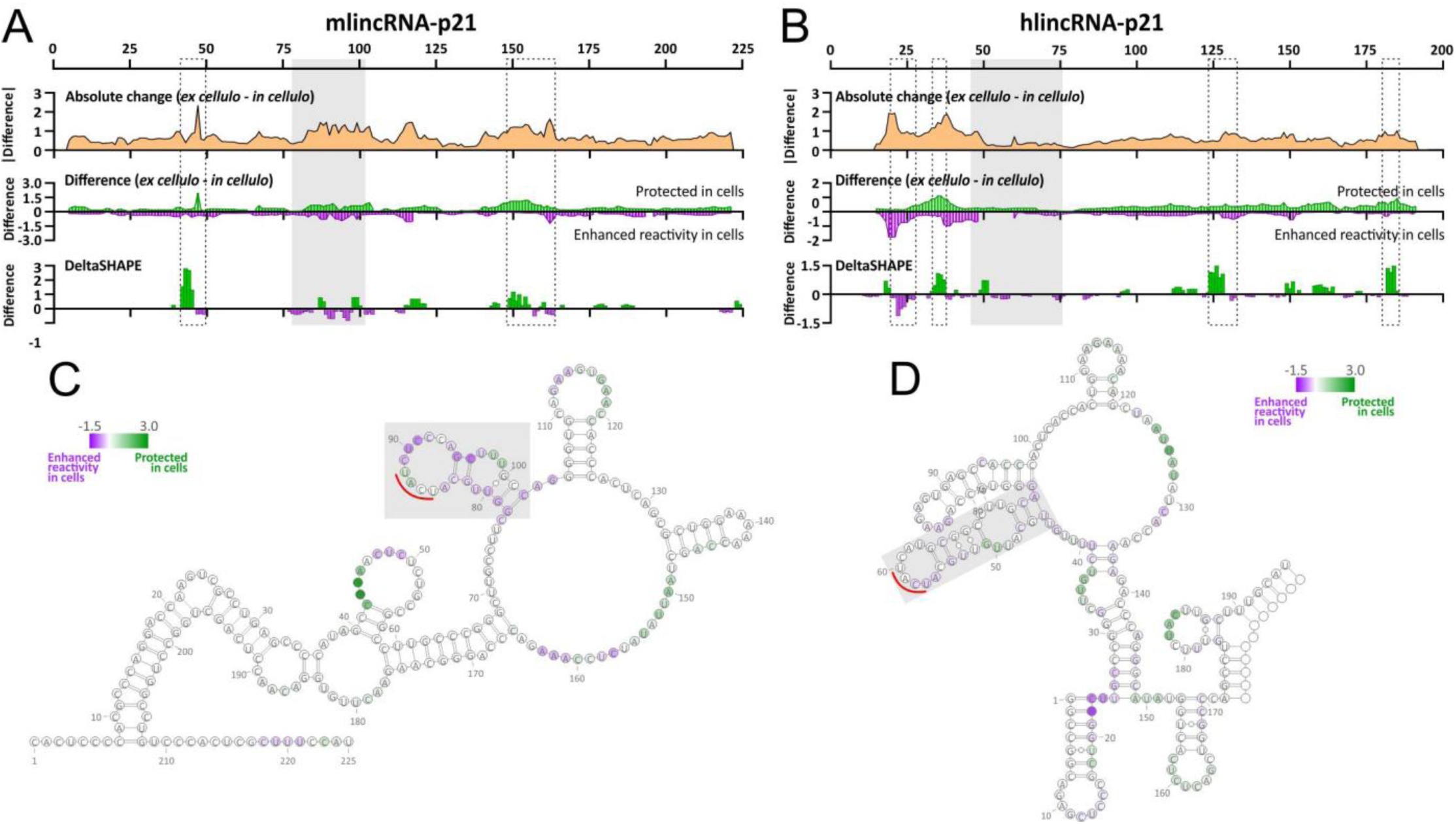
The comparison of *ex cellulo* and *in cellulo* SHAPE-MaP profiles indicates protein binding to the putative hnRNP K-binding site on lincRNA-p21. A and B. ΔSHAPE analyses for mlincRNA-p21. (**A**) and hlincRNA-p21 (**B**). (Top panel) The absolute change in SHAPE reactivity between *ex cellulo* and *in cellulo* states, smoothed over 9-nt windows. (Middle panel) Contributions of positive and negative differences (green and violet, respectively) to the absolute change, where positive values indicate protection in cells, and negative values indicate enhanced reactivity in cells due to structural reorganization, smoothed over 9-nt windows. (Lower panel). ΔSHAPE values indicating regions of the RNA exhibiting statistically significant changes between *in cellulo* and *ex cellulo* conditions. The grey box represents the position of the putative hnRNP K helical arm. The dotted boxes represent other statistically-significant regions of protein binding. C and **D.** Schematic representation of the 5’-end secondary structures of the mlincRNA-p21 (**C**) and hlincRNA-p21 (**D**), gradient color-coded at single-nucleotide resolution representing the ΔSHAPE values. The grey box represents the position of the putative hnRNP K helical arm.

In summary, our ΔSHAPE analysis showed consistent recognition of specific structural domains by endogenous proteins in both RNAs, including the putative hnRNP K-binding structure.

### hnRNP K binds lincRNA-p21 through its UCAU-containing helical arm inside cells

To further test whether the UCAU-containing helical arm mediates hnRNP K binding in cells, we performed eCLIP-qRT-PCR. We first confirmed antibody specificity for endogenous hnRNP K by Western blotting (Fig. S9A). We then transfected murine NIH-3T3 cells with a plasmid containing either the wild-type 3-kb-long full-length mlincRNA-p21 construct (WT-FL-mlincRNA-p21) or a mutated version lacking the UCAU-containing helical arm (mut-FL-mlincRNA-p21). HCT116 cells received the corresponding 4-kb hlincRNA-p21 constructs (WT-FL-hlincRNA-p21 or mut-FL-hlincRNA-p21). WT transcripts showed slight enrichment relative to mutants (Fig. 7A, S9B), but the differences were not statistically significant. To rule out nonspecific protein binding in the very long FL constructs, we repeated the assay with shorter constructs (Fig. 5A-B). The WT:mutant median ratio improved (mouse: 1.32 to 2.24; human: 1.11 to 2.22) but remained non-significant, with overlapping individual values (Fig. 7B, right).

**Figure 7.**
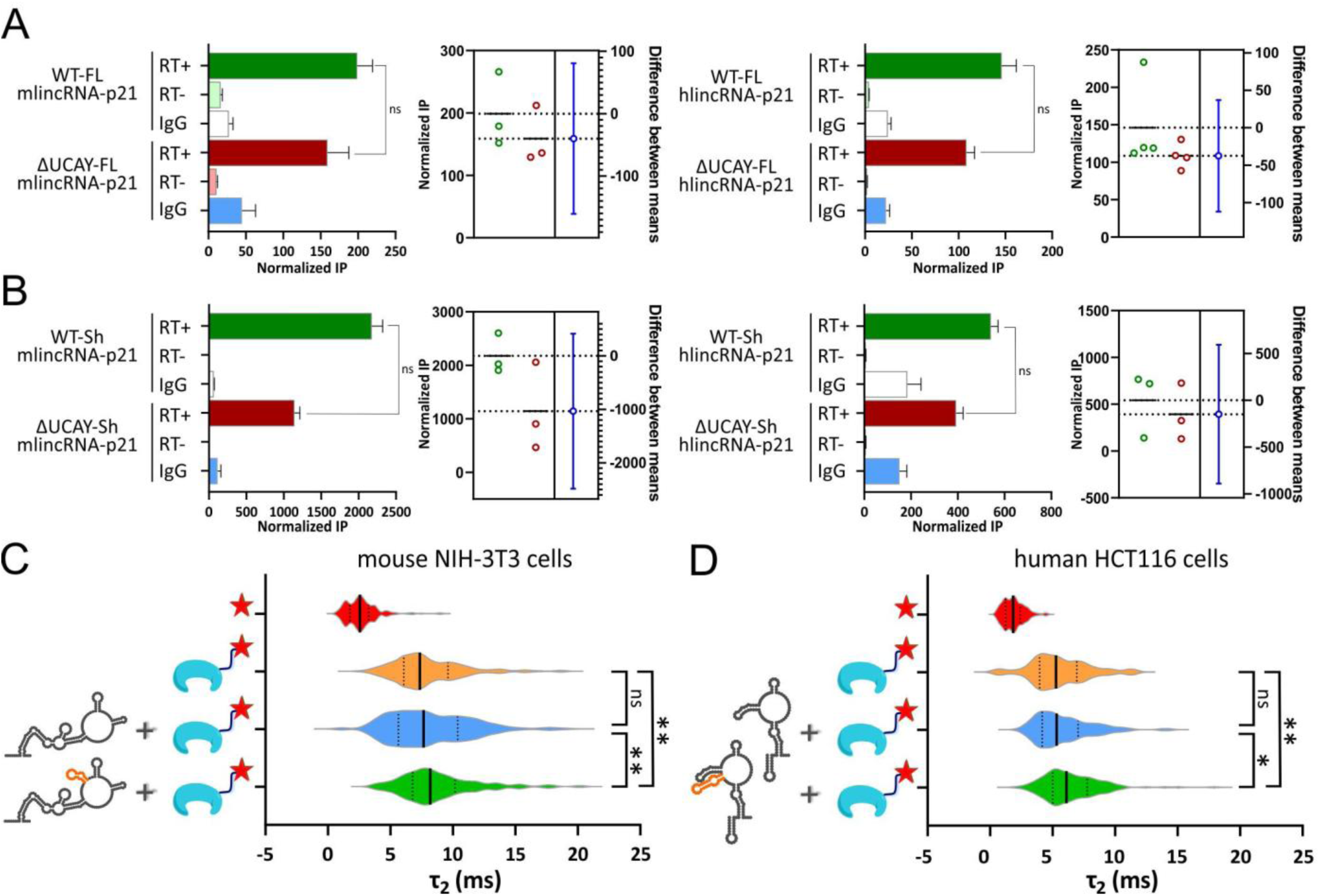
The UCAU-containing helical arm in lincRNA-p21 is bound by hnRNP K inside cells. A and. **B.** Normalized immunoprecipitation (IP) results from the eCLIP-qRT-PCR assay using full-length (FL) (A) or short-constructs (B) of mlincRNA-p21 (left panels) and hlincRNA-p21 (right panels) constructs. The IP was normalized against the input expression values (Fig. S9), the nonspecific U6 transcript as a loading control, and the neomycin transcript as a transfection control. RT-indicates % of IP-RT-with IP-WT, not enrichment with input. Error bars represent the calculated propagation error of 3 biological replicates. Estimation plots showing the difference between means are displayed on the right side of each graph, indicating the precision of the calculated effect size as a 95% confidence interval. **C and D.** Fluorescence Correlation Spectroscopy (FCS) assay with violin plots representing the correlation time associated with the bound species (τ_2_) for each transfected system in mouse (C) and human (D) cells. The τ_2_ value represents the average time it takes for the fluorescent molecule to traverse the confocal detection volume in the bound or complexed state. From top to bottom, red indicates cells transfected with 10 pmol of plasmid encoding the fluorescent tdTomato protein, and 10 pmol of an empty pcDNA3 plasmid; orange indicates cells transfected with 10 pmol of the fusion hnRNP K-tdTomato protein, and 10 pmol of an empty pcDNA3 plasmid; green indicates cells transfected with 10 pmol of the fusion hnRNP K-tdTomato protein, and 10 pmol of the WT pcDNA3-lincRNA-p21 short construct; blue indicates cells transfected with 10 pmol of the fusion hnRNP K-tdTomato protein, and 10 pmol of the mutant pcDNA3-lincRNA-p21 short construct. Asterisks represent statistically significant differences (P < 0.05) as determined by lognormal Brown-Forsythe and Welch ANOVA tests.

Given these narrow differences, we reasoned that bulk, population-averaged eCLIP-qRT-PCR might mask discrete, low-occupancy binding events. We therefore used live-cell Fluorescence Correlation Spectroscopy (FCS), an assay that detects individual or few-molecule binding events by measuring the residence time of a fluorescently tagged molecule in the confocal volume. Here, we measured changes in hnRNP K residence time upon binding of lincRNA-p21, either WT or a mutant lacking the UCAU-containing helical arms. Co-transfecting fluorescently tagged hnRNP K with untagged WT or mutant short constructs (Fig. 7C-D) showed significantly longer hnRNP K residence times with WT than mutant RNA, in both mouse and human cells (Fig. 7C-D), indicating a higher bound fraction and supporting hnRNP K binding *via* the UCAU-containing helical arm. As observed in our cell-based assays (ΔSHAPE and eCLIP-RT-qPCR), the results showed replicate-to-replicate variation (Fig. S9). Within individual replicates, the broader range of residence times observed for the hnRNP K fusion samples (orange, green, blue) compared to the narrow tdTomato control (red) may reflect cell-to-cell heterogeneity in the fraction of hnRNP K engaged with lincRNA-p21 relative to its other physiological targets, adding a further layer of biological complexity ^31^.

In summary, our *in cellulo* binding assays support the proposal that both mlincRNA-p21 and hlincRNA-p21 bind hnRNP K through the helical arm containing the UCAU tetranucleotide, in murine and human cells, respectively. The variability observed both between replicates and within individual cell populations is consistent with a low-affinity, non-canonical binding interaction whose occupancy is sensitive to cellular context, rather than a fixed, high-affinity interaction.

## DISCUSSION

In this work, we applied a combined approach to demonstrate that two syntenic lncRNAs, mouse and human lincRNA-p21, which diverge overall in sequence and structure, establish a conserved RNA-protein interaction with hnRNP K. Using *in silico*, *in vitro*, *ex cellulo*, and *in cellulo* methods, we show that both mlincRNA-p21 and hlincRNA-p21 transcripts contain a structural motif comprising a conserved UCAU tetranucleotide that binds the KH3 RNA-binding domain of hnRNP K (Fig. 8A). Deletion of this motif reduces the interaction between lincRNA-p21 and hnRNP K both in mouse and human cells. Our approach, which is based on the general paradigm that conserved sequence and structure reflect conserved biological function, suggests that the mechanism by which lincRNA-p21 activates *Cdkn1a* transcription through hnRNP K binding, so far demonstrated only in mice, may also extend to humans, and potentially to other mammals, and offers a general strategy to discover shared structural elements in other poorly conserved lncRNAs.

**Figure 8.**
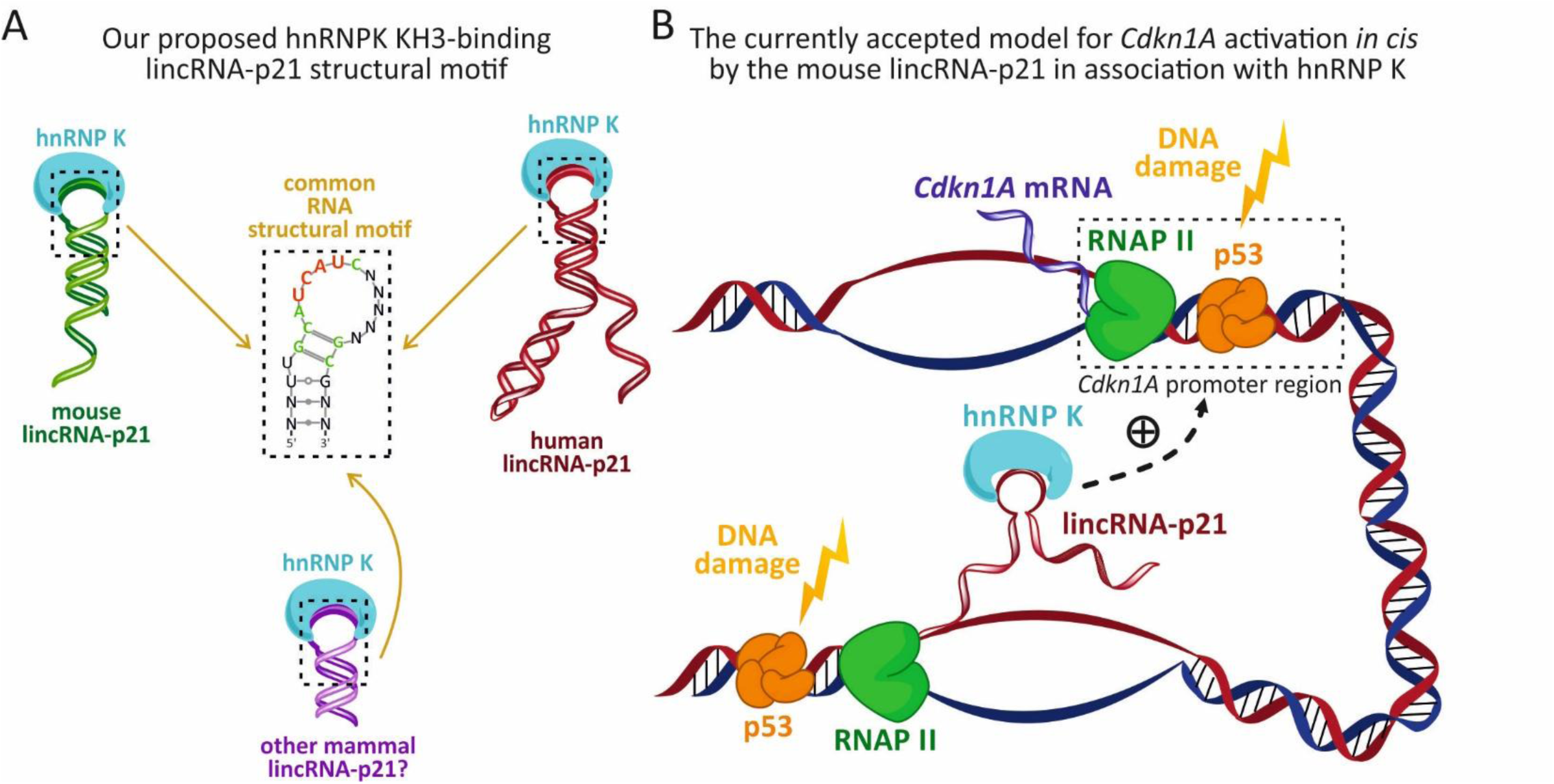
The structural elements of the conserved interaction between lincRNA-p21 and the hnRNP K protein in the p21 DNA damage pathway. **A**. A conserved RNA structural motif binds the KH3 domain of the hnRNP K protein. A helical arm containing the hairpin with the essential UCAU tetranucleotide (red) and conserved residues (green) is found in the mlincRNA-p21 and hlincRNA-p21, and other mammal lincRNA-p21 transcripts. The KH3 domain of the hnRNP K protein binds the helical arm containing this RNA motif in mouse and human cells, and evolutionary conservation suggests a broader distribution in other mammalian species. **B**. Currently accepted model for the mouse lincRNA-p21-dependent activation of the *Cdkn1A* gene locus in association with the hnRNP K protein, in agreement with our proposed hnRNP K binding structural motif. LincRNA-p21 activates the transcription of the *Cdkn1A* gene *in cis*. The nascent lincRNA-p21 transcript, which is physically tethered to the locus, forms the UCAU-containing helical arm co-transcriptionally, leading to the recruitment of the hnRNP K protein through the interaction of its KH3 domain. The lincRNA-p21-hnRNP K complex can localize in physical proximity to the *Cdkn1A* locus, acting as a coactivator for p53-dependent *Cdkn1A* mRNA transcription, as previously proposed ^9,10^.

### A combined structural and functional approach uncovers a conserved interaction between lincRNA-p21 and hnRNP K

The poor sequence conservation of lncRNAs necessitates alternative strategies for identifying functional elements. Syntenic conservation alone cannot resolve fast-evolving, sequence-degenerate genes ^32^. Also, k-mer screening, while useful for detecting functional convergence, lacks resolution where sequence itself is not conserved ^33^. Structural approaches, including covariance models that align structure rather than sequence, offer a complementary route ^34^. Given the caveats of each method, we combined sequence alignment, structural covariation, *in silico* analyses, molecular dynamics simulations, and experimental (*in vitro*, *ex cellulo*, and *in cellulo*) assays to identify a functional element shared between mouse and human.

This approach identified a helical arm within a four-way junction, comprising a core UCAU tetranucleotide conserved across >100 mammals, that binds the KH3 domain of hnRNP K, similar to a SELEX-derived NOVA-2 KH3 domain-binding element ^25,26^. Yet, this interaction appears weaker than the SELEX-identified one. Both lncRNA orthologs lack the conserved ACCNNN hairpin-loop sequence beyond the essential UCAY tetranucleotide, and the hydrogen bond between the second-position cytidine and Glu-14 (mutated to Lys in hnRNP K) is absent. We disagree, however, with a previously published proposal that a third-position adenosine sterically clashes with hnRNP K Arg-40 ^26^. Structural superposition shows the relevant hydrogen bond distance (2.2 Å) is shorter than the reported clash-generating distance (3.2-3.6 Å) (Fig. S10A-B). Moreover, despite good UCAC/dTCCC superposition (RMSD 0.505 Å), nucleoside geometry differences allow minor interface rearrangements (Fig. S10C), consistent with the structures’ resolution (1.8-2.4 Å) ^35^. Indeed, the UCCC-to-UCAC mutation reduced affinity (from Kd ∼2 to ∼50 µM) without abolishing binding, as shown by NMR ^26^. Moreover, our molecular dynamics simulations (MDS) support this steric compatibility, and show that the third adenosine forms stable hydrogen bonds with Ile-49. Also consistent with our molecular dynamics simulations, complementary *in vitro* binding assays showed that both orthologs’ 5’ structures bound full-length hnRNP K and its KH3 domain, but not KH1 or KH2.

Inside cells, our eCLIP-qRT-PCR showed no significant WT-mutant difference, despite a visible WT enrichment. This is relevant in light of other examples of weak RNA-protein interactions, such as those involving the PRC2 complex. PRC2 components and hnRNP K both lack protein domains with a high propensity for UV crosslinking, as is the case for RRM RNA-binding domains ^36^. Lack of statistically significant differences and replicate variability can also reflect the weak nature of the lincRNA-p21 and hnRNP K interaction, which shows variable RNA signal occupancy and a crosslink yield proportional to RNA occupancy on the protein ^36^. Indeed, an *in cellulo* technique not subjected to these limitations, such as live-cell single (or few)-molecule Fluorescence Correlation Spectroscopy (FCS), revealed significant WT-mutant differences in hnRNP K binding. However, binding was incomplete at any given moment, consistent with a genuinely weak interaction.

Together, these results indicate that mlincRNA-p21 and hlincRNA-p21 bind hnRNP K’s KH3 domain through a weak, transient interaction mediated by a conserved, non-canonical UCAU-containing motif (Fig. 8A), a property we propose reflects the functional requirement for a reversible interaction that depends on the cellular stress state (see below).

### Significance of the lincRNA-p21 interaction with hnRNP K for gene expression during the DNA damage stress response

MlincRNA-p21 has been proposed to act as a coactivator that recruits hnRNP K and p53 to the *Cdkn1A* promoter *in cis* ^9^. Specifically, exon 1 of nascent mlincRNA-p21 was shown to contain regulatory elements required for *Cdkn1a* activation ^10^. A mutant spanning position 76-262 (and thus, containing the hnRNP K-binding helical arm) significantly reduced *Cdkn1a* expression, whereas a mutant spanning the exon1-intron splice junction (and thus, leaving the helical arm intact) had no effect ^10^. These results, consistent with our finding that the hnRNP K-binding motif lies at the 5’ end, support the current model in which nascent mlincRNA-p21, physically tethered to the gene, acts as a local recruitment platform (Fig. 8B).

HlincRNA-p21, in contrast, has been reported to have functions distinct from mlincRNA-p21, including cytoplasmic translation inhibition and hypoxia-enhanced glycolysis ^37,38^, indicating functional divergence. Although hlincRNA-p21 was known to bind hnRNP K ^11^, whether it activates *Cdkn1a* transcription remained unclear. Given our combined evidence (sequence conservation and structural similarity, *in vitro* and *in cellulo* binding, *in silico* stability), we propose that this mlincRNA-p21-dependent *Cdkn1A*-activation mechanism extends to hlincRNA-p21.

We further show that the hnRNP K-binding element comprises a non-canonical UCAU-containing motif, conserved in 101/111 analyzed mammals. Though it diverges from the consensus U/CCCC or CCCCCC sequence ^39,40^, it was previously identified *via* three-hybrid screening ^41^ and is compatible with the CNCNCNCNNNCC pattern from RNA Bind-n-Seq and eCLIP data ^42^. This contrasts with SELEX data showing that the CCCC-to-CCAC mutation increases dissociation by 3.5-fold ^43^. This discrepancy likely reflects assay differences, since Bind-n-Seq captures sequence enrichment rather than highest-affinity motifs. Notably, high affinity is not always evolutionarily favored. Many eukaryotic transcription factors instead achieve specificity through locally elevated concentration ^44^, achievable *via* transcriptional hubs or phase separation, which is consistent with our previous finding that hlincRNA-p21 colocalizes with nuclear paraspeckles ^8^.

We propose that this weak and dynamic UCAU-hnRNP K KH3 interaction is essential for finely tuned *Cdkn1a* expression during the DNA damage response, where sustained p21 accumulation could impair homeostasis recovery. This contrasts with hnRNP K’s interaction with XIST via a poly-C motif, which instead requires strong, sustained binding to maintain durable X-chromosome inactivation ^39,45^.

## CONCLUSIONS

In this work, we show that two syntenic lncRNAs with globally divergent sequences and structures can establish a conserved RNA-protein interaction via a shared structural motif, a finding that would not have been possible using just one technique. Only by combining sequence alignment, structural covariation, molecular dynamics, and orthogonal binding assays across murine and human cellular contexts could we resolve a non-canonical RNA motif, comprising a conserved UCAU sequence within the helical arm of a shared four-way junction at the 5’-end of lincRNA-p21. This motif links mlincRNA-p21 and hlincRNA-p21 to the same protein, hnRNP K, through the same structural logic. This strongly suggests that hlincRNA-p21 activates *Cdkn1a* transcription by the same mechanism long described only in mice. If this extension holds true, it will close a gap that has persisted in the field for over a decade.

This RNA motif may have remained hidden due to its lower affinity for KH3 than the canonical poly-C-rich motif. More broadly, the weakness of this interaction is itself the finding. A tight, stable RNA-protein complex would lock *Cdkn1a* in an “on” state; a weak, transient one would allow the cell to respond to DNA damage and then return to its basal state. This alleged discrepancy highlights the plasticity of evolution in acquiring new genetic elements while preserving others, thereby promoting adaptation to changing genetic environments.

## MATERIALS AND METHODS

### Cloning and mutagenesis

The vector pcDNA3-WT-FL-mlincRNAp21 was a kind gift of Maite Huarte (CIMA, University of Navarra, Spain) ^7^. The vector pcDNA3-WT-FL-hlincRNAp21 was previously cloned from the cDNA of the human HEK-293 cell line ^8^. Using these vectors as templates, the vector pcDNA3-ΔUCAY-FL-mlincRNA-p21 was created through a two-step overlapping PCR amplification, first using the primers 120_F_979_QC_del-hnRNPK (GCTTGCCTTCGGTTGCTTTGCCAGGGG) with pcDNA3_Rev (GATCAGCGAGCTCTAGCA) and pcDNA3-F-Amp (ACTGTCATGCCATCCGTAAG) with 121_R_1018_QC_del-hnRNPK (CCCCTGGCAAAGCAACCGAAGGCAAGC), followed by a second PCR with the primers pcDNA3-Fwd (CCACTGCTTACTGGCTTATC) and pcDNA3-Rev. The resulting product was digested with the restriction enzymes *Kpn*I and *Eco*RI and ligated into a previously digested vector pcDNA3-WT-FL-mlincRNAp21, which lacked the WT insert. The vector pcDNA3-ΔUCAY-FL-hlincRNA-p21 was also created through a two-step overlapping PCR amplification. First, the primers 111_F_QC_dHNRNPKmotif (GGGCTTGTCTTTTGTTGCATTGCCTTGCAGGGTGCCAGAA) with 26_R_1000_hInt (CCAGAGCTAAAAGGATGTGAAGC) and pcDNA3-F-Amp with 112_R_QC_dHNRNPKmotif (TTCTGGCACCCTGCAAGGCAATGCAACAAAAGACAAGCCC) were used. This was followed by a PCR using the external primers pcDNA3-F-Amp and 26_R_1000_hInt. The resulting product was digested with the restriction enzymes *Srf*I and *Nhe*I and ligated into a previously digested vector, pcDNA3-WT-FL-hlincRNAp21, which lacked the WT insert. The short lincRNA-p21 constructs were made by amplifying the 5’-end region of the pcDNA3-WT-FL-mlincRNAp21, pcDNA3-ΔUCAY-FL-mlincRNAp21, pcDNA3-WT-FL-hlincRNAp21, and pcDNA3-ΔUCAY-FL-hlincRNAp21 vectors. To create the pcDNA3-WT-Sh-mlincRNA-p21 and pcDNA3-ΔUCAY-FL-mlincRNAp21 vectors, a 553 bp fragment was amplified with primers 08F_CMVprom (CCTACTTGGCAGTACATCTACG) and 125_R_243_EcoRI-mLIncp21 (GGGGAATTCGGACAGGCCAGGCCAG), containing an *Eco*RI restriction site. To create the pcDNA3-WT-Sh-hlincRNA-p21 and pcDNA3-ΔUCAY-FL-hlincRNAp21 vectors, a 485 bp fragment was amplified with primers 08F_CMVprom and 124_R_152_EcoRI-hLIncp21 (GGGGAATTCATATGCCCTGGGTCTCTTG), also containing an *Eco*RI restriction site. All PCR fragments were further digested with the restriction enzymes *Kpn*I and *Eco*RI and inserted into pcDNA3-WT-FL-mlincRNAp21 or pcDNA3-WT-FL-hlincRNAp21, which have been previously digested with the same enzymes and lacked the FL inserts. To express the hnRNP K protein and its KH1, KH2, and KH3 domains, the parental vector pT7-V5-SBP-C1-HshnRNPK, which was a gift from Elisa Izaurralde (Addgene plasmid # 64923; http://n2t.net/addgene:64923; RRID: Addgene_64923), was used for PCR amplification. The FL hnRNP K fragment was amplified using primers 01_F_pET_FL (CCCCCATGGGAATGGAAACTGAACAGCCAGA) and 02_R_pET_FL (GGGGGTACCTTAGAATCCTTCAACATCTGCAT), the KH1 domain, the primers 03_F_pET_KH1 (CCCCCATGGGAATGGTTGAATTACGCATTCTGC) and 04_R_pET_KH1 (GGGGGTACCTTAGATTTTCTTCAGAATTTCTCCAATTG), the KH2 domain, the primers 05_F_pET_KH2 (CCCCCATGGGAGACTGCGAGTTGAGGCTG) and 06_R_pET_KH2 (GGGGGTACCTTAGATGATCTTTATGCACTCTACAACC), and the KH3 domain, the primers 07_F_pET_KH3 (CCCCCATGGGAATTATTACTACACAAGTAACTATTCCC) and 08_R_pET_KH3 (GGGGGTACCTTACAGCAAATACTGTGCATTCTG). The hnRNP K FL amplicon was digested with the restriction enzymes *Nco*I and *Kpn*I and inserted into an empty, previously digested pETM44 plasmid with the same restriction enzymes to create a pETM44-hnRNPK-FL vector. The hnRNP K KH1, KH2, and KH3 amplicons were digested with the restriction enzymes *Nco*I and *Kpn*I and cloned into a pET-MBP-EGFP vector, previously digested with the same restriction enzymes to create pET-MBP-hnRNPK-KH1, pET-MBP-hnRNPK-KH2, and pET-MBP-hnRNPK-KH3. For the FCS assays, the pcDNA3-tdTomato and pcDNA3-hnRNPK-tdTomato vectors were created using the pT7-V5-SBP-C1-HshnRNPK plasmid as a template for the hnRNP K gene, and the Zurkowska_pBSK_TomBlast_pLann plasmid, a kind gift from Dr. Maud Borensztein (Institute of Molecular Genetics of Montpellier, France), as a template for the tdTomato gene. To create the pcDNA3-hnRNPK-tdTomato vector, the hnRNPK gene was amplified from the pT7-V5-SBP-C1-HshnRNPK plasmid using the primers 09_F_NheI_K_hnRNPK (GGTGCTAGCGCCACCATGGAAACTGAACAGCCAGAAG) and 11_R_hnRNPK_3KpnI (GGGGTACCACTGAATCCTTCAACATCTGCATACTGCT) and was further digested with the restriction enzymes *Nhe*I/*Kpn*I. The tdTomato gene was amplified from the Zurkowska_pBSK_TomBlast_pLann plasmid with the primers 04_F_tdTomato_5KpnI (TTTGGTACCTCTGTGAGCAAGGGCGAGGA) and 02_R_tdTomato_3EcoRI (GCGCGCGAATTCTTACTTGTACAGCTCGTCCA) and was further digested with the restriction enzymes *Nhe*I/*Eco*RI. To create the pcDNA3-tdTomato vector, the tdTomato gene was amplified from the Zurkowska_pBSK_TomBlast_pLann plasmid using the primers 01_F_tdTomato_5NheI (ATTGCTAGCGCCACCATGGTGAGCAAGGGCGAG) and 02_R_tdTomato_3EcoRI, and was further digested with the restriction enzymes *Kpn*I/*Eco*RI. To accommodate all inserts, the empty pcDNA3 plasmid was amplified using primers pcDNA3_R_NheI_SLIC (GGCGCTAGCAATTTCGATAAGCCAGTAAGCAG) and pcDNA3_F_EcoRI_SLI (CCGAATTCCAGCCTCGACTGTGCCTT), and then further digested with the restriction enzymes *Kpn*I/*Eco*RI. The pcDNA3 *Kpn*I/*Eco*RI digested plasmid was ligated to the hnRNPK *Kpn*I/*Nhe*I and the tdTomato *Nhe*I/*Eco*RI fragments to create the pcDNA3-hnRNPK-tdTomato vector, and to the tdTomato *Kpn*I/*Eco*RI fragment to create the pcDNA3-tdTomato vector.

### *In vitro* transcription and purification of mlincRNA-p21 and hlincRNA-p21

The mlincRNA-p21 and hlincRNA-p21transcripts were produced and purified under non-denaturing conditions, as previously described ^8,13^ with minor modifications. Briefly, the plasmids pcDNA3-WT-FL-mlincRNAp21 and pcDNA3-WT-FL-hlincRNA-p21 were linearized overnight with the restriction enzyme *Eco*RI (NEB) and purified with a G-25 spin column (Cytiva). The linearized vector was transcribed *in vitro* using T7 polymerase in 1X transcription buffer containing 40 mM Tris-HCl (pH 8.0), 8 mM MgCl2, 100 mM KCl, 5 mM DTT, 0.01% Triton X-100, and 0.2 mg/ml molecular biology-grade BSA (NEB). Following transcription, the template DNA and proteins were removed with 25 U of Turbo DNase (Thermo Scientific) per ml of transcription and 0.2 mg/ml of proteinase K (Euromedex), respectively. The transcribed RNA was rebuffered in MOPS filtration buffer (8 mM MOPS-K buffer pH 6.5 and 100 mM KCl) using Amicon Ultra-0.5 centrifugal concentrators (molecular weight cut-off of 100 kDa) (Millipore). The RNA was further purified by size-exclusion chromatography (SEC) using Tricorn columns (GE Healthcare) self-packed with Sephacryl S1000 resin (Cytiva) and run in 0.1 M KCl, 8 mM K-MOPS pH 6.5, 0.1 mM Na-EDTA.

### *In vitro* Selective 2’-Hydroxyl Acylation analyzed by Primer Extension (SHAPE)

SHAPE analysis was performed according to Chillon, et al. ^13^. Briefly, primers containing an amine terminal group were labeled with 5-FAM (5-Carboxyfluorescein, succinimidyl ester) (tebu-bio) and JOE (6-Carboxy-4’,5’-dichloro-2’,7’-dimethoxyfluorescein, succinimidyl ester) (tebu-bio) (440-660 ng/μl of oligonucleotide, 2.5 μg/μl SE-dye freshly dissolved in DMSO and 75 mM sodium tetraborate buffer pH 8.5) in an overnight reaction at room temperature. Labeled primers were purified in a denaturing 20% polyacrylamide gel and dissolved in TE buffer (100 mM Tris-HCl pH 8.5, 10 mM EDTA-Na pH 8.5). A sequencing ladder was generated in a reaction using primers labeled with 5-FAM and a solution containing 1.6 μl of a 70 ng/µl hLincRNA-p21 LIsoE2 plasmid, 1.84 μl of Thermo Sequenase buffer (Affymetrix), 4 μl of 2 μM FAM-coupled primer, 16 μl of ddT mix (150 μM dATP, 150 μM dCTP, 150 μM dGTP, 150 μM dTTP, 1.5 μM ddTTP), 1.84 μl of Thermo Sequenase DNA Polymerase (Affymetrix), and 7.52 μl of water. PCR was run for 40 cycles using an annealing temperature of 55 °C for 20 s, and an extension time of 1 min at 72 °C. The PCR product was precipitated, and the pellet was dissolved in 40 μl of formamide. *In vitro* transcribed hlincRNA-p21 was purified under native conditions and folded in the presence of 5 mM magnesium chloride and a monovalent ion mixture (200 mM KCl, 50 mM Na-HEPES pH 7.4, and 0.1 mM EDTA-Na pH 8.5) for 30 min at 37°C. The SHAPE reaction was performed by incubating 30 pmol of RNA with 2.5 mM 1-methyl-7-nitroisatoic anhydride (1M7) (synthesized by the Chemical Core Facility at EMBL Heidelberg), 2.5 mM N-methylisatoic anhydride (NMIA) (Sigma), 2.5mM 1-methyl-6-nitroisatoic anhydride (1M6) (Sigma), and an equal volume of dimethyl sulfoxide (DMSO) as a negative control in a final volume of 0.4 ml and 5 min of incubation time at 37°C, in duplicate ^14,15^. Chemically modified RNA was reverse transcribed using 24 primers, positioned every 200 bp of the hlincRNA-p21 (Table S1). 1 pmol of RNA was annealed to JOE-coupled primers (0.1 μM primer, 0.1 mM EDTA-Na pH 8.5, 1 M betaine) in a final volume of 12 μl, heated at 95°C for 2 min, and cooled to 48°C. Reverse transcription was initiated by adding 8 μl of a mixture containing 2.5X reverse transcriptase buffer (Thermo), 12.6 mM DTT, 944 mM betaine, 1.26 mM dNTP mix, and 120 U Superscript III reverse transcriptase (Thermo). The resulting DNA was precipitated, and the pellet dissolved in 40 μl of formamide. Samples and ladders were loaded in a 96-well plate [3 μl of M7-treated or DMSO-treated sample, 4 μl DNA ladder, and 14 μl formamide]. Samples were then submitted for fragment length analysis with capillary electrophoresis (Eurofins). QuShape ^46^ was used to determine the chemical probing reactivity profiles. Formation of adducts was quantified by comparison between the 1M7-, 1M6-, NMIA-, and the DMSO-treated samples. Average values of individual 1M7 reactivity values from 2 experiments were normalized with the “simple2boxplot.py” Python script ^15^, and average values of individual 1M6 and NMIA reactivity values from 2 experiments were normalized with “boxplot2simple.py”Ppython script ^15^. Typical reads from consecutive primers overlapped by about ∼20 nucleotides. In these overlapping regions, we averaged reactivity values from the two contributing primers in each replicate before averaging the corresponding values of independent replicates. Normalized 1M6 reactivity values were subtracted from the NMIA reactivity values with the “differenceByWindow.py” Python script ^15^. The software SuperFold with default settings was used to obtain the secondary structure maps ^47^.

### *In vitro* Selective 2’-hydroxyl acylation analyzed by primer extension and mutational profiling (SHAPE-MaP)

SHAPE-MaP analysis was performed according to Smola, et al. ^16^. *In vitro* transcribed mlincRNA-p21 was purified under native conditions and folded in the presence of 5 mM magnesium chloride and a monovalent ion mixture (200 mM KCl, 50 mM Na-HEPES pH 7.4, and 0.1 mM EDTA-Na pH 8.5) for 30 min at 37°C. The SHAPE reaction was performed by incubating 15 pmol of RNA with 10 mM 1-methyl-7-nitroisatoic anhydride (1M7) (synthesized by the Chemical Core Facility at EMBL Heidelberg), 10 mM N-methylisatoic anhydride (NMIA) (Sigma), 10 mM 1-methyl-6-nitroisatoic anhydride (1M6) (Sigma), and an equal volume of dimethyl sulfoxide (DMSO) as a negative control in a final volume of 0.04 ml and 15 min of incubation time at 37°C, in triplicate. RNA was precipitated with 100 % ethanol and washed with both 100 % and 70 % ethanol before resuspending it in 20 µl of water. Sites of chemical modifications in the RNA were converted into mutations using error-prone reverse transcription as in Mustoe et al (2019) ^48^. Briefly, 1 pmol of mlincRNA-p21 chemically-modified RNA was incubated with 200 ng of random nonamers (Thermo) and 2 mM dNTPs at 65 °C for 10 min followed by 4 °C for 2 min. Nine uL of 2.22X MaP buffer (6 mM MnCl_2_, 1 M betaine, 50 mM Tris pH 8.0, 75 mM KCl, 10 mM DTT) and 1 μL of SuperScript II Reverse Transcriptase (Invitrogen) were added to the reaction and the combined solution incubated at 23 °C for 2 min followed by: 25 °C for 10 min, 42 °C for 90 min, 10 X [50 °C for 2 min, 42 °C for 2 min], and 72 °C for 10 min. The resulting cDNA was purified with the QIAquick PCR & Gel Cleanup Kit (Qiagen) and used as a template for a PCR reaction. The mlincRNA-p21 was covered using 6 overlapping PCR reactions (Table S2). The PCR products were cleaned with the Monarch® PCR & DNA Cleanup Kit (5 μg). Sequencing libraries were created using the Nextera XT kit (Illumina) according to the manufacturer’s protocol and sequenced on an Illumina MiSeq-250 PE at the EMBL Genomics Core Facility (Genecore). Mutations in sequencing reads were converted into reactivity profiles using ShapeMapper 2 ^49^. Average values of 1M7, 1M6, and NMIA were obtained from three experiments. Averaged 1M6 reactivity values were subtracted from the averaged NMIA reactivity values with the “differenceByWindowSHAPEMAP.py” Python script to get the differential reactivity values ^16^. RNA secondary structures were modeled using the folding pipeline Superfold ^16^ with the 1M7, and the differential reactivity values, serving as constraints for minimum free energy calculations. Secondary structure maps were visualized using the Java Applet VARNA ^50^.

### In cellulo SHAPE-Map

*In cellulo* SHAPE-Map was performed according to Smola, et al. ^30^ and Smola and Weeks ^51^ with modifications. *In cellulo* SHAPE-MaP reactivity values were obtained for the mlincRNA-p21 and the hlincRNA-p21 after transfecting 1250 fmol of plasmids into NIH-3T3 mouse and HCT116 human cells, respectively, seeded in 10-cm cell culture plates. Cells were crosslinked with 4000 μJ/cm^2^ of 254 nm UV light, dislodged with a cell scraper, and spun down by centrifugation at 450 g for 5 min at 4 °C. The pellet was resuspended in 1 ml of ice-cold lysis buffer [40 mM of Tris-HCl pH 8, 25 mM of NaCl, 6 mM of MgCl_2_, 1 mM of CaCl_2_, 256 mM of sucrose, 0.5 % of Triton X-100, 0.5 U/μl of RNAsin (Promega), and 0.45 U/μl of DNaseI (Roche)], and incubated at 4°C for 5 min. The nuclei were pelleted by centrifuging at 1500 g at 4°C for 4 min. Pellets were resuspended in 0.95 ml of PBS at room temperature and split into two tubes by placing 0.45 ml into each tube. To each tube, 50 µL of 2M 2-methylnicotinic acid (NAI) (synthesized by the Chemical Core Facility at EMBL Heidelberg) (C_F_ = 200 mM) or DMSO was added, and the tubes were incubated for 15 min at room temperature with rotation. Nuclei were collected by centrifuging at 1500 g for 5 min at 4°C, and the pellet resuspended in 100 μl of proteinase K digestion buffer (40 mM of Tris-HCl pH 8, 200 mM of NaCl, 1.5 % of SDS, and 500 μg / ml of Proteinase K (Euromedex). Tubes were incubated with the proteinase K for 45 min at room temperature, with continuous rotation. The RNA was extracted with the Monarch RNA cleaning kit-5 (NEB) and eluted in 83 μl of water. DNA was eliminated by further incubation with 5 µl of TURBO DNase (2 U/μl) (Thermo), 2 μl Murine RNase H inhibitor (40 U/µl) (NEB) in a final volume of 100 µl. The RNA was purified with the Monarch RNA cleaning kit-5 (NEB) and eluted in 30 μl of water. Mutational profiling (MaP) reverse transcription (RT) was performed as in Mustoe, et al. ^48^ with minor modifications. Briefly, 3.5 µg of nuclear RNA was mixed with 200 ng of random nonamers (NEB) and 10 mM of dNTP in a volume of 10 µl, and incubated at 65 °C for 10 min followed by 4 °C for 2 min. 9 uL 2.22x MaP buffer [1x MaP buffer consists of 6 mM MnCl_2_, 1 M betaine, 50 mM Tris (pH 8.0), 75 mM KCl, 10 mM DTT] was added and the combined solution was incubated at 23 °C for 2 min. Finally, 1 μL SuperScript II Reverse Transcriptase (Thermo) was added and the RT reaction was performed according to the following temperature program: 25 °C for 10 min, 42 °C for 90 min, 10 × [50 °C for 2 min, 42 °C for 2 min], 72 °C for 10 min. The reaction was stopped by heating at 72°C for 10 min. The cDNA was purified with the PCR CleanUP kit (Qiagen) and eluted in 30 μl. The cDNA was further amplified by PCR using the Q5 hot start DNA Polymerase (NEB) and the primers pcDNA3_F_5’MCS (TACCGAGCTCGGATCCACT) and 02_R_648_Exon2 (GTGTCAATGCTCTCGCTATG) to produce an amplicon of 738 bp of the mlincRNA-p21, or the primers 92_F_2_Start (GCCGGCAGAGCTCC) and 87_R_799_hEx1 (AGGCTGTATGCACTTTATTACCT) to produce an amplicon of 795 bp of the hlincRNA-p21. The PCR products were purified with the Monarch® PCR & DNA Cleanup Kit (5 μg) (NEB) and eluted in 15 μl. Sequencing libraries were created using the Nextera XT kit (Illumina) according to the manufacturer’s protocol and sequenced on a MiSeq (EMBL GeneCore) or DNBSEQ-G400 instrument (BGI) with 250-PE or 150-PE, respectively. Mutations in sequencing reads were converted into reactivity profiles using ShapeMapper 2 (Busan and Weeks 2018). Reactivity values were used in the calculation of the deltaSHAPE values (see below).

### Ex cellulo SHAPE-Map

*Ex cellulo* SHAPE-Map was performed according to Smola, et al. ^30^ and Smola and Weeks ^51^ with modifications. *Ex cellulo* SHAPE-MaP reactivity values were obtained for the mlincRNA-p21 and the hlincRNA-p21 after transfecting 1250 fmol of plasmids into NIH-3T3 mouse and HCT116 human cells, respectively, seeded in 10-cm cell culture plates. The cells were dislodged with Trypsin-EDTA (0.05%) (Thermo), and spun down by centrifugation at 450 g for 5 min at 4 °C. The pellet was resuspended in 1-2 ml of ice-cold lysis buffer [40 mM of Tris-HCl pH 8, 25 mM of NaCl, 6 mM of MgCl_2_, 1 mM of CaCl_2_, 256 mM of sucrose, 0.5 % of Triton X-100, 0.5 U/μl of RNAsin (Promega), and 0.45 U/μl of DNaseI (Roche)], and incubated at 4°C for 5 min. The nuclei were pelleted by centrifuging at 1500 g at 4°C for 4 min. The pellet was resuspended in 0.5 ml of proteinase K digestion buffer (40 mM of Tris-HCl pH 8, 200 mM of NaCl, 1.5 % of SDS, and 500 μg / ml of Proteinase K (Euromedex) and incubated at room temperature for 45 minutes. The RNA was extracted twice with phenol/chloroform/isoamyl alcohol solvent (24:24:1), pre-equilibrated with 1.1× folding buffer (111 mM of HEPES pH 8, 165 mM of KCl, and 5.55 mM of MgCl_2_), and once with chloroform. Buffer exchange was performed with 1.1× folding buffer over an Amicon 0.5-ml 100K cut-off (Millipore), spinning four times at 5000 g for 3 min each. After the cleaning, the RNA was equilibrated at 37°C for 20 min. The RNA was then split into two tubes, each one containing 0.45 ml. To each tube, 50 µL of 22M -methylnicotinic acid (NAI) (synthesized by the Chemical Core Facility at EMBL Heidelberg) (C_F_ = 200 mM) or DMSO was added, and the tubes were incubated for 15 min at room temperature with rotation. The RNA was extracted with the Monarch RNA cleaning kit-5 (NEB) and eluted in 83 μl of water. DNA was eliminated by further incubation with 5 µl of TURBO DNase (2 U/μl) (Thermo), 2 μl Murine RNase H inhibitor (40 U/µl) (NEB) in a final volume of 100 µl. The RNA was purified with the Monarch RNA cleaning kit-5 (NEB) and eluted in 30 μl of water. Mutational profiling (MaP), reverse transcription (RT), library preparation, sequencing, and sequence analysis were performed as for *in cellulo* SHAPE-MaP.

### Differences in SHAPE reactivity (Delta-SHAPE)

Delta-SHAPE was performed as described in Smola, et al. ^30^. Briefly, the reactivity values of the *ex cellulo* SHAPE-MaP, showing high correlation, were averaged. The *in cellulo* SHAPE-MaP values were not averaged, and a delta-SHAPE analysis was performed for each *in cellulo* SHAPE-MaP dataset compared with the averaged *ex cellulo* SHAPE-MaP dataset. To assess consistent protein binding sites among replicates, the averaged *ex cellulo* and *in cellulo* SHAPE-MaP datasets were also analyzed. To obtain the difference (*ex cellulo – in cellulo*), the reactivity values of the *in cellulo* SHAPE-MaP were subtracted from those of *ex cellulo* SHAPE-MaP, and smoothed over a 9-nt window. To obtain the absolute change *(ex cellulo – in cellulo),* the subtracted values were converted into their absolute values and smoothed over a 9-nt window. The statistically significant differential regions (DeltaSHAPE) were calculated using the Python script “deltaSHAPE.py” ^30^.

### Molecular dynamics simulations of free lincRNA-p21

Molecular dynamics simulations were performed on the stems of the helical arm containing the UCAU tetranucleotide. The starting coordinates for free RNA were obtained using the web server of the simRNA software package ^27,28^, where we imposed the experimental secondary structure obtained by SHAPE and SHAPE-MaP, and performed the Monte Carlo generation procedure with preset parameters. The coordinates were then processed in the software tleap to hydrate and neutralize the system, and subsequently create topology and configuration files suitable for simulations with AMBER^29^. The molecular mechanics model for simulations of conformational energies and noncovalent interactions Parm99 was considered ^52^ with the bsc0+χOL3 modifications ^53,54^. Water was modeled according to the TIP3P model ^55^, while system neutralization was achieved by adding a suitable number of sodium atoms described by the Joung-Cheatham parameterization ^56^. The cutoff for van der Waals interactions and short-range electrostatics was set to 9 Å, while long-range electrostatics was accounted for by particle-mesh Ewald. Simulations were performed in AMBER with the pmemd.cuda routine optimized for GPU parallelization. A standard simulation protocol was performed, consisting of four steps: minimization, thermalization, equilibration, and production. In the minimization step, the system underwent a minimization procedure to prevent steric clashes. To achieve this, only water molecules and ions were adapted by applying a strong positional restraint to the RNA, and considering 2500 steepest-descent steps, followed by an additional 2500 steps with the conjugate gradient method. Then, the same minimization process was repeated without restraints. After minimization, the system was slowly thermalized from 0 to 300 K in a constant-volume simulation of 300 ps. The temperature was set according to a Langevin thermostat with a collision frequency of 1 ps-1 and a time step of 2 fs. Bonds involving hydrogens were constrained using the SHAKE algorithm. In the equilibration step, the system was allowed to adjust its density through a 20 ns simulation at a constant pressure of 1 bar, imposed by a Berendsen barostat. Finally, a 1μs NVT simulation was performed for production purposes, where the coordinates of the system were saved every 20 ps. Throughout the simulation, a restraint was applied to the last base pair of the helical arm (G-78:C-102 and U-45:A-74 for mlincRNA-p21 and hlincRNA-p21, respectively) to compensate for the lack of stability introduced by the absence of the rest of the molecule. To this end, the NMR optimization protocol of the AMBER suite was employed, which introduced suitable distance and angle restraints for the base pair, with strengths of 1.44 kcal·mol-1·Å-2 and 1.44 kcal·mol-1, respectively.

### Molecular dynamics simulations of the lincRNA-p21/KH3 complex

The starting coordinates for the simulations of the complex were obtained by considering the hnRNP K KH3 domain in complex with a 6-mer ssDNA as reported in the PDB structure 1ZZI ^26^. After discarding three external nucleotides, the protein in complex with the central dTCCC tetramer was obtained. Then, the trajectory of the free lincRNA-p21 was aligned to best match the backbone of UCAU to the coordinates of dTCCC within the complex. The snapshot with the lowest root-mean-square displacement was selected, and a coordinate file, containing the KH3 domain and the aligned UCAU tetramer, was created. To model the free RNA, the coordinates were fed to tleap for preprocessing, and the system was minimized, equilibrated, and simulated according to the procedure above, although the production step was limited to 500 ns. To model the protein, the ff14SB model was employed ^57^. The last snapshot of this preliminary simulation was used as a template for preparing the lincRNA-p21/KH3 complex. A protocol involving two main steps was used: free RNA adjustment and complex relaxation. In the first step, the last snapshot of the free RNA simulation was aligned with the template to match the coordinates of the UCAU tetramer. The resulting coordinates were used to prepare the input files with tleap and minimize the system’s energy. Next, a series of 1 ns NVT simulations was performed, where the UCAU atoms were restrained to the template coordinates with increasing strength, ranging from 0.001 to 10 kcal·mol-1·Å-2. To avoid perturbing the secondary structure of the RNA, distance and angle restraints were added for each base pair with the same parameters as above. The final coordinates of the RNA were then combined with the KH3 template coordinates and fed to tleap for the complex relaxation step. To avoid steric clashes between RNA and the protein, due to the large size of lincRNA-p21, a soft-core version of the van der Waals interaction was employed ^58^. Specifically, a first imposition of λ=0 for the RNA fragments outside the binding was used, which turned off their van der Waals interactions with the rest of the system. With this setup, short simulations were performed where the RNA molecule was gently steered to disentangle it from the protein. The presence of positional restraints on the tetramer and distance/angle restraints on the base pairs allowed the maintenance of the RNA secondary structure and the binding interface of the complex during the process. After this, further 0.5 ns NPT simulations in which λ was increased in a stepwise manner were performed. The system was then equilibrated in a 20 ns NPT simulation, during which the van der Waals interactions regained their full strength. At the same time, the restraint on the tetramer coordinates was removed while the secondary-structure restraints remained in place. Finally, the system was simulated in a 1μs NVT run where only the end base pair was restrained, as in the case of free RNA.

### Analysis of Molecular Dynamics simulations

The overall convergence of the simulations was assessed by monitoring the time evolution of the root-mean square displacement (RMSD) (Fig. S4D). Hydrogen bonds and van der Waals contacts were detected according to geometric criteria based on the routines “hbond” and “nativecontacts” of the software CPPTRAJ ^59^. Threshold values of 3.5 Å and 135° for hydrogen bonds and 4.0 Å for van der Waals contacts were applied. In the case of secondary structures, a base pair was considered to be formed when all the relevant hydrogen bonds were present simultaneously. Errors in the figures were computed by combining bootstrap and block averaging ^60^.

### Conservation analysis

The genomes of 111 mammal species used in the structure-based conservation analysis were chosen according to the trees provided by the UCSC Genome Browser (https://github.com/ucscGenomeBrowser/kent/blob/master/src/hg/utils/phyloTrees/213way.comm <u>onNames.nh</u>) (downloading information summarized in Table S3). The downloaded .fasta sequences were used to build a local BLAST database with the makeblastdb tool of the BLAST+ package (NCBI). The custom-made BLAST database was searched against the cDNA sequence of mlincRNA-p21 and hlincRNA-p21 using the tool blastn (dc-megablast option). Extracted sequences were manually curated to eliminate redundancy. Manually curated sequences were aligned using the MuscleWS tool within the Jalview program ^61,62^. To analyze structure-based conservation, a covariation model was created using the mlincRNA-p21 Superfold-derived secondary structure and 42 seed sequences that showed high sequence similarity and a lack of gaps with the cmsearch tool from the Infernal software package^63^. The remaining mammalian sequences were aligned to the lincRNA-p21 covariance model using the Infernal tool cmalign. To assess statistically significant conserved RNA structural motifs, lincRNA-p21 secondary structure covariation was measured using R-scape ^64^ with the Lancaster covariation aggregation option ^65^.

### *In vitro* transcription with Cy5-UTP labeling

For *in vitro* transcription with Cy5-UTP labeling, the plasmids producing the short lincRNA-p21 constructs, pcDNA3-WT-Sh-mlincRNA-p21 and pcDNA3-WT-Sh-hlincRNA-p21, were used. To generate the DNA templates for transcription, plasmids were linearized at the 3’ end of the lncRNA sequence using the *Eco*RI restriction site. Internally labelled RNAs were produced by *in vitro* transcription using the HighYield T7 RNA Labelling Kit (Jena Bioscience) using UTP-X-Cy5 nucleotides as previously described ^66^. Each reaction contained 500 ng of DNA template, 0.2 μl of Cy5-UTP (2.5 mM), and 0.4 μl of RiboLock RNase Inhibitor (Thermo), and was incubated for 4 hours at 37°C. The DNA template was digested with 1 μl TURBO™ DNase (Thermo) for 15 minutes at 37°C. Labelled RNA probes were purified using ProbeQuant™ G-50 Micro Columns (Cytiva) and eluted in 50 μl.

### *In vitro* protein-RNA UV-crosslinking assay

His-tagged proteins were produced from BL-21 (RIPL) bacterial strain, purified on Ni-NTA agarose, eluted from the beads with Imidazole, and quantified by Coomassie staining as described in Carnesecchi, et al. ^66^. The assay was performed as previously described ^66^. Briefly, binding assays were performed in a volume of 30 μl containing 1x binding buffer (20 mM HEPES pH 7.9, 1.4 mM MgCl2, 1 mM ZnSO4, 40 mM KCl, 0.1 mM EDTA, 5% Glycerol), 2 μg tRNA (Thermo), 3 μg BSA, 10 mM DTT and 0.1% NP40. 2 pmol of internally-UTP-Cy5-labelled lincRNA-p21 probes were added with 0.5-1 µg of *in vitro* purified hnRNPK FL protein, or the isolated KH1, KH2 or KH3 domains. After 20 min on ice, the samples were irradiated with UV light (UVP-Crosslinker, Jena Analytik) for 10 min and transferred to Eppendorf tubes. 0.8 µl of RNase A (Thermo) was added, and the samples were incubated for 20 min at 37°C. Cy5-labelled RNA-protein complexes were resolved on 10-12% SDS-PAGE for 40 minutes at 180 V and detected by fluorescence (imager IQ800 Cytiva). Following the detection, the gels were stained with Coomassie overnight, rinsed with water, imaged, and quantified using the Fiji software.

### Enhanced crosslinking and immunoprecipitation coupled to reverse transcription and quantitative PCR (eCLIP-RT-qPCR)

For the eCLIP-RT-qPCR, the anti-hnRNP K polyclonal antibody (RN019P, MBL) was chosen from the ENCODE database for antibody selection (https://www.encodeproject.org/search/?type=AntibodyLot&status=released) and the procedure followed the eCLIP protocol described previously with modifications ^67^. Human HCT116 and mouse 3T3 cells were seeded on two 10-cm plates and transfected with 217.32 fmol/plate of each of the lincRNA-p21 constructs tested using Lipofectamine 2000 (Thermo). After 36 hours, cells were crosslinked with 254 nm UV light at 4000 x 100 µJ/cm^2^. Cells were trypsinized, spun down for 5 min at 200 g, and resuspended in 1 ml of PBS. 1 ml of iCLIP lysis buffer [50 mM of Tris-HCl pH 7.4, 100 mM of NaCl, 1 % of Igepal CA630, 0.1 % of SDS, 0.5 % of sodium deoxycholate, 5.5 µl of 1:200 Protease inhibitor Cocktail III (Calbiochem) per 1 ml iCLIP lysis buffer, 11 µl murine RNase inhibitor (NEB) per 1 ml of lysis buffer] was added per sample (if 20 million cells or adjusted depending on the actual cell number). Cells were incubated on ice for 15 min. In the meantime, magnetic Dynabeads M-280 sheep anti-rabbit (Thermo) were washed twice with 500 µl of cold iCLIP lysis buffer. For 20 million cells, 10 µg of antibody was added to 100 µl of washed beads, and incubated at room temperature for 45 min. After lysis, cells were added 30 µl of TURBO DNase (Thermo), incubated at 37°C for 30 min. Debris spun down at 15000 g, 4°C for 5 min, and the supernatant was transferred to new Eppendorf tubes. The beads were further washed with 500 µl of cold iCLIP lysis buffer and resuspended in 100 µl of cold iCLIP lysis buffer. Add the washed beads to 20 million cells. Incubate horizontally at 4°C for 2 hours. After the incubation, 15 % of the sample was set aside as input and treated with 200 µl of Proteinase K mix [100 mM of Tris-HCl pH 7.4, 50 mM of NaCl, 10 mM of EDTA-Na pH 8, 25 µl (20 mg/ml, Euromedex)] and incubated for 20 min at 37°C with rotation at 1200 rpm. RNA was extracted with 0.75 ml of Trizol LS reagent (Thermo). After RNA extraction, the 83 µl of RNA were treated with 5 µl of TURBO DNase (40 U/µl, Thermo) in a final volume of 100 µl, and incubated at 37°C for one hour. The RNA was further purified with the Monarch RNA cleaning kit-5 (NEB) and eluted in 30 μl of water. The rest of the sample (85 %) was washed twice with 0.9 ml of cold High salt wash buffer (50 mM of Tris-HCl pH 7.4, 1 M of NaCl, 1 mM of EDTA-Na pH 8, 1 % of Igepal CA630, 0.1 % SDS, and 0.5 % of sodium deoxycholate), followed by two washes with 0.5 ml of wash buffer (20 mM of Tris-HCl pH 7.4, 10 mM of MgCl_2_, and 0.2 % of Tween-20) using a magnetic stand. The sample was finally resuspended in 85 µl of wash buffer, 15 µl (15 % of the initial amount) set aside for Western blot analysis, and the rest (70 % of the initial amount), treated with Proteinase K as described above and RNA extracted (IP sample). The input and IP RNA was converted into cDNA by adding 200 ng of random primers (Thermo) and 10 mM of dNTP. The sample was heated at 65°C for 5 min on a thermocycler and cooled down to 25°C. 200 units of SuperScript™ IV reverse transcriptase (Thermo) were added, and the mixture was incubated at 25°C for 10 min, followed by 50°C for 2 hours. At the end of the reaction, RNA-DNA duplexes were eliminated with 0.5 µl of *E.coli* RNase H (NEB) by incubating at 37°C for 30 min. The mouse and human lincRNA-p21 cDNA was amplified by qPCR using the primers 122_F_49_mLincp21 (CCCATAGCCACAACTCTCTG) and 123_R_146_mLincp21 (CTGAGTGGGTGGTTCACTTC) for the mlincRNA-p21, and 92_F_2_Start (GCCGGCAGAGCTCC) and 53_R_96_hEx1 (GGCTCACTCTTCTGGCAC), which spans the expected hnRNP K binding site. Additionally, all samples were amplified for the neomycin gene, with primers pcDNA3-F-Neo (TGGATTGCACGCAGGTTCT) and pcDNA3-R1-Neo (GGACAGGTCGGTCTTGACA), which is expressed from the pcDNA plasmid, as a control for transfection efficiency, and for the housekeeping gene U6 with primers U6_Fwd (CTTCGGCAGCACATATACTAA) and U6_Rev (AATATGGAACGCTTCACGAAT), as a control for sample loading.

### Western blotting

A total of 2 % from the input and 15 % of the IP samples from the eCLIP-RT-qPCR assay were set aside for Western blot analysis. To these samples, 7.5 μl of 4X Laemmli buffer (Bio-Rad) was added and cold iCLIP lysis buffer up to 20 μl. Before boiling the sample at 95°C for 5 min, 3 µl of 1M DTT was added, and 15 µl of sample were loaded into a 4-12 % Bis-Tril gel, 10-well, 1.5 mm (Thermo), using MES SDS running buffer (Thermo). The gel was run at 100 V for 5 min, followed by 200 V for 40 min, and the proteins were transferred to a Immun-Blot PVDF membrane (Bio-Rad) using a wet-transfer Blotting System (Bio-Rad) using 100 V for 1 hour. The membrane was blocked with TBST buffer (10 mM Tris-HCl pH 8, 150 mM of NaCl, and 0.05 % (w/v) of Tween20) and 5% BSA at room temperature for 1 hour. A mixture of primary antibodies, including 1:200 of Anti β-actin (Santa Cruz Biotechnologies, sc-47778) and 1:1000 of Anti-hnRNP K (MBL, RN019P), was used, and the mixture was incubated with the membrane at room temperature for 1 hour. The membrane was washed 3 times with TBST buffer for 5 min each, and incubated with a mixture of secondary antibodies: 1:1000 Goat anti-mouse IgG (H+L)– Alexa647 (Thermo) and 1:1000 Goat anti-rabbit IgG (H+L)– Alexa488 (Thermo), and incubated at room temperature for 1 hour. The membrane was washed three times with TBST buffer and a final time with PBS, before visualizing it on a ChemiDoc Imaging System (Bio-Rad).

### Fluorescence Correlation Spectroscopy (FCS)

Human HCT116 (38200 cells) and mouse 3T3 (25500 cells) cell lines were seeded on Nunc Lab-Tek II Chambered Coverglass 2-wells (Thermo). For the tdTomato alone condition, 20 fmol of pcDNA3-tdTomato and 20 fmol of pcDNA3-empty plasmids were transfected into each cell line using Lipofectamine 2000. For the hnRNPK-tdTomato condition, 20 fmol of pcDNA3-tdTomato and 20 fmol of pcDNA3-hnRNPK-tdTomato plasmids were transfected. For the lincRNA-p21 WT condition, 20 fmol of pcDNA3-hnRNPK-tdTomato and 20 fmol of pcDNA3-WT-Sh-mlincRNA-p21 or pcDNA3-WT-Sh-hlincRNA-p21 were transfected. For the mutant condition, 20 fmol of pcDNA3-hnRNPK-tdTomato and 20 fmol of pcDNA3-ΔUCAY-Sh-mlincRNAp21 or pcDNA3-ΔUCAY-Sh-hlincRNAp21 were transfected. The FCS data were acquired using a 40X C-APO water 1.4NA on a Confocal Zeiss LSM980 NLO located at the Montpellier Ressources Imagerie (MRI) light microscopy facility. For each cell and specified point, 10 measurements of 5 s each were taken at 0.1 % of 561 nm laser power. The data were fit to a Confocal (Gaussian): 3D + 3D model using the PyCorrFit software (https://github.com/FCS-analysis/PyCorrFit/releases/tag/1.1.7) and the Levenberg-Marquardt algorithm with no weights. The values of the correlation time associated with the bound species (τ2) and its associated error were extracted for each measurement. To eliminate outliers, the percentage of error was calculated for each measurement, and values exceeding 25% were removed.

## ACKNOWLEDGMENTS

We thank Robert Feil and Anna Marie Pyle for their critical reading of the manuscript. We also thank members of Feil’s team and Odil Porrua for their constructive input. IC is grateful to the funds provided by the *Fondation pour la recherche sur le cancer*, ARC (ARCPJA2021060003686), and *La Ligue Contre le Cancer Pyrénées-Orientales* 2023 and 2025. This work was partly funded by ITMO Cancer (18CN047-00, awarded to MM) and by the Fondation ARC pour la recherche sur le cancer (PJA-20191209284, awarded to MM). MM is also partly funded by the Swedish National Research Council (VR, 2024-04107), Cancerfonden (project 25 4360 Pj), SciLifeLab (Strategic initiatives and technology development call 2025), and the Italian Association for Cancer Research (AIRC, IG 28746). SA acknowledges support from a Ramón y Cajal Fellowship (ref. RYC2022-037744-I), funded by MICIU/AEI/10.13039/501100011033 and FSE+, and from the Spanish Ministerio de Ciencia e Innovación (MCIN) through the project PID2023-149150OB-I00. JC acknowledges the AFM Téléthon (Trampoline grant ID 24140), which supported the recruitment of CB.

## AUTHOR CONTRIBUTIONS

SA designed and performed the Molecular Dynamics Simulations. CB performed the *in vitro* protein-RNA UV-crosslinking assay. CF designed the Fluorescence Correlation Spectroscopy assay. JC designed and quantified the *in vitro* protein-RNA UV-crosslinking assay. MM obtained funding and contributed to the analysis and interpretation of the results. IC conceived, designed, and supervised the project, obtained funding, performed and analyzed all other assays (*in vitro*, *in cellulo*, and *ex cellulo* SHAPE and SHAPE-MaP, eCLIP-RT-qPCP, and FCS experiments, and the *in silico* analysis of crystal structures), and wrote the manuscript with input from all authors.

---

**Figure S1.**
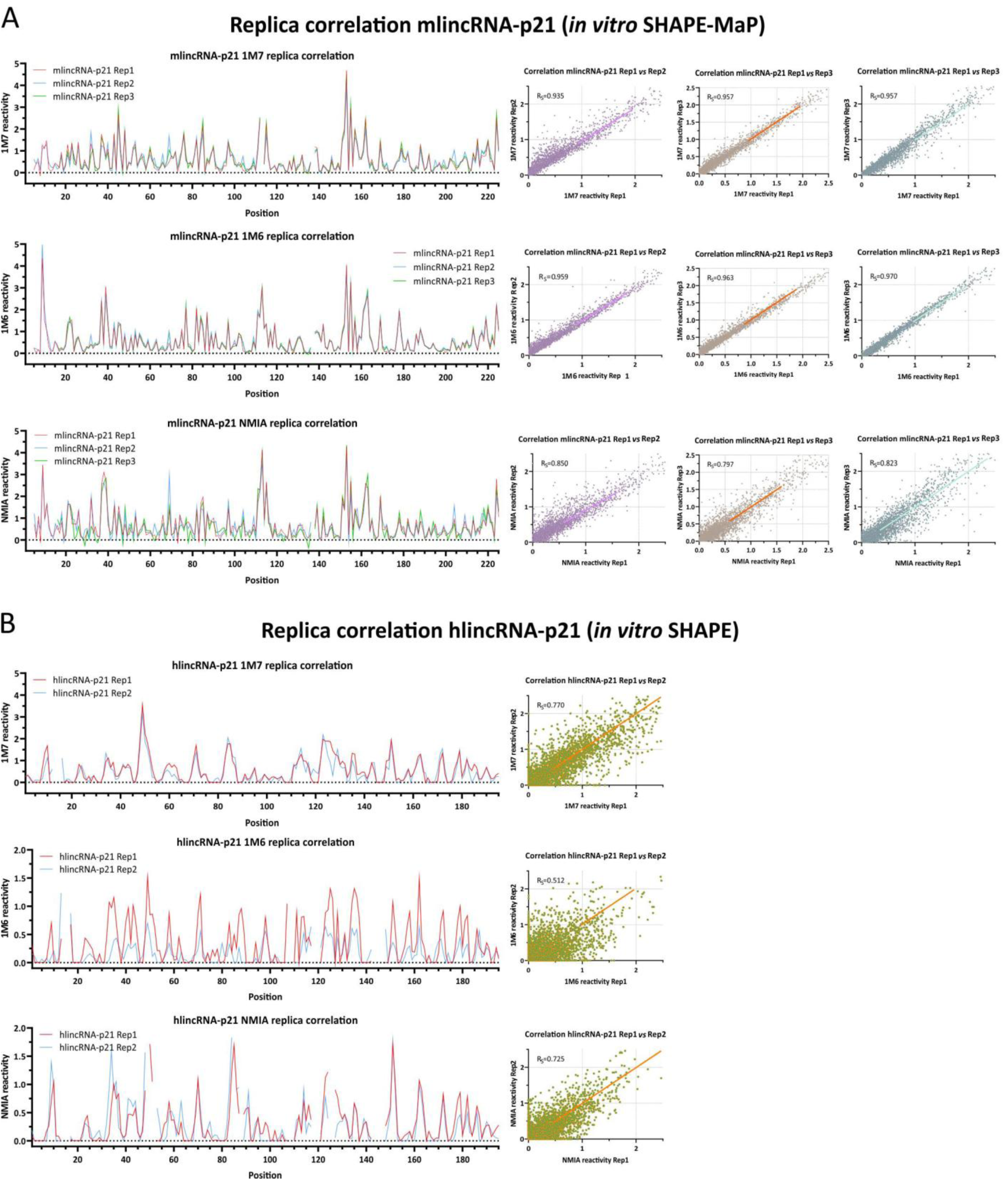
Replicate correlation for SHAPE and SHAPE-MaP analyses (related to. Figure 1). (Left panels) Reactivity values obtained after treatment with 1M7, 1M6, and NMIA reagents for each individual replicate of mlincRNA-p21 **(A)** and hlincRNA-p21 **(B).** (Right panels) Scatter plots representing the reactivity values for each pair of replicates. Spearman correlation coefficients are indicated for each comparison.

**Figure S2.**
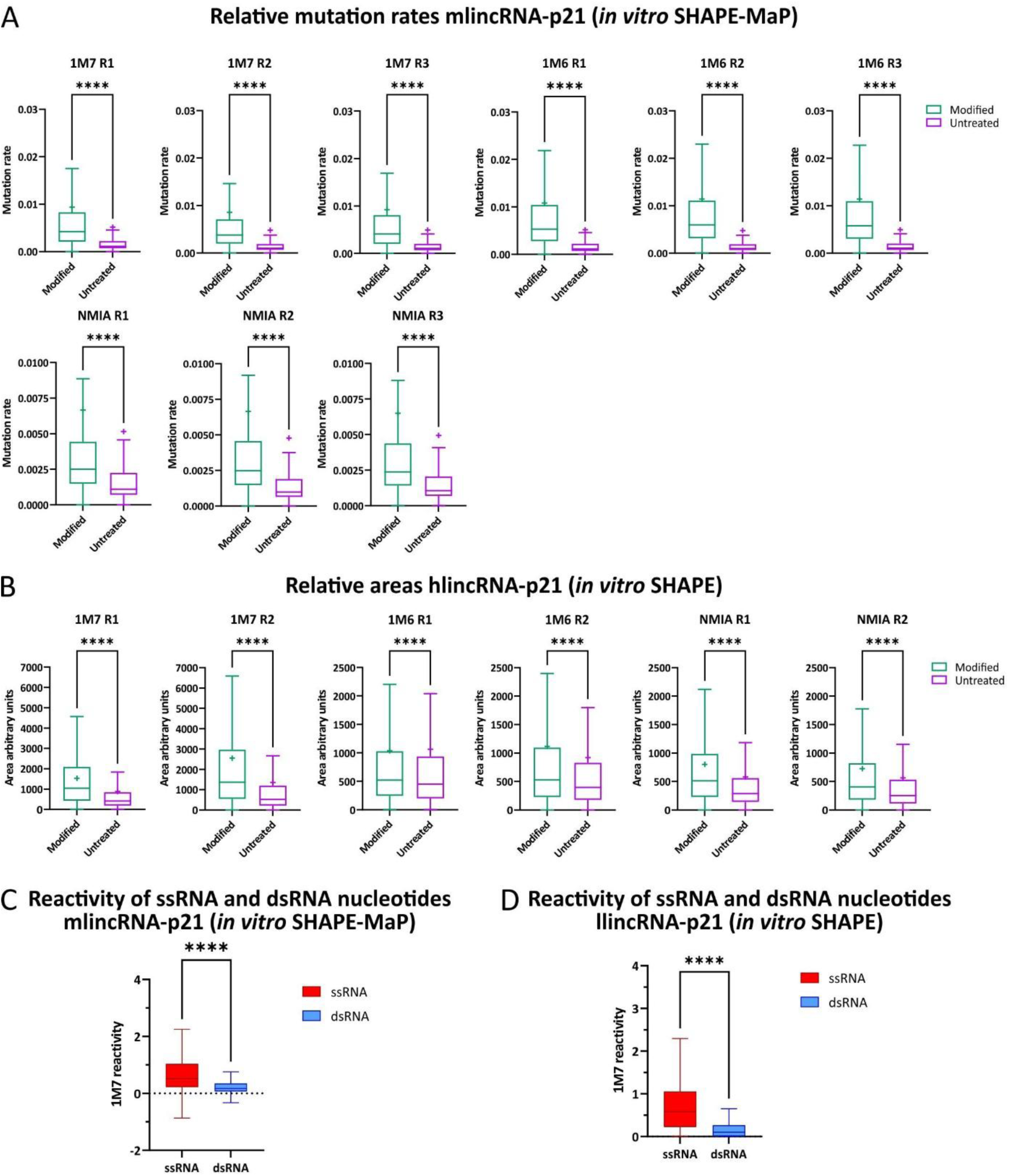
Quality control metrics for the *in vitro* SHAPE and SHAPE-MaP (related to. Figures 1 **and 2). A and B.** Mutation rates for the three biological replicas of mlincRNA-p21 *in vitro* SHAPE-MaP (A) and two replicas of hlincRNA-p21 *in vitro* SHAPE (B) (box, interquartile range [IQR]; median indicated by line; average indicated by “x”; whiskers are drawn in the Tukey style, and values outside this range are not shown). **C and D.** Mapping of 1M7 reactivity data to single- and double-stranded regions for the first 225 nt of the mlincRNA-p21 (C) and first 195 nt of the hlincRNA-p21 (D) (box, interquartile range [IQR]; median indicated by line; whiskers are drawn in the Tukey style, and values outside this range are not shown), corresponding to the secondary structures depicted in Fig. 2A.

**Figure S3.**
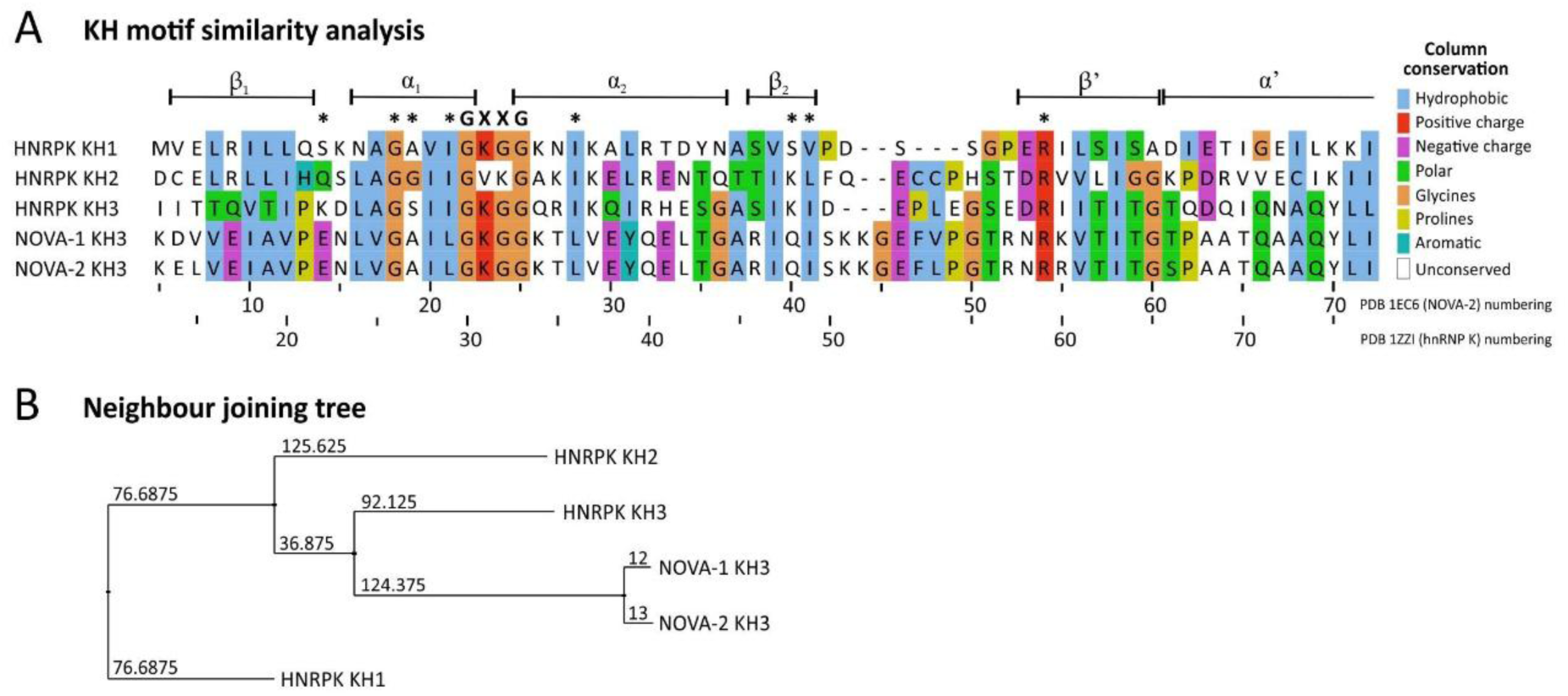
KH3 domain sequence and structure comparison (related to. Figure 2**). A.** Sequence alignments of the three KH domains of hnRNP K and the KH3 domains of NOVA-1 and NOVA-2 proteins, obtained by TCoffeeWS software. GXXG represents the conserved loop of the DNA-binding cleft. Asterisks indicate RNA-contacting residues. Numbering according to the PDB 1EC6 structure**. B.** Neighbor-joining tree obtained from the Clustal X alignment in the Jalview program. Values indicate estimated phylogenetic distances.

**Figure S4.**
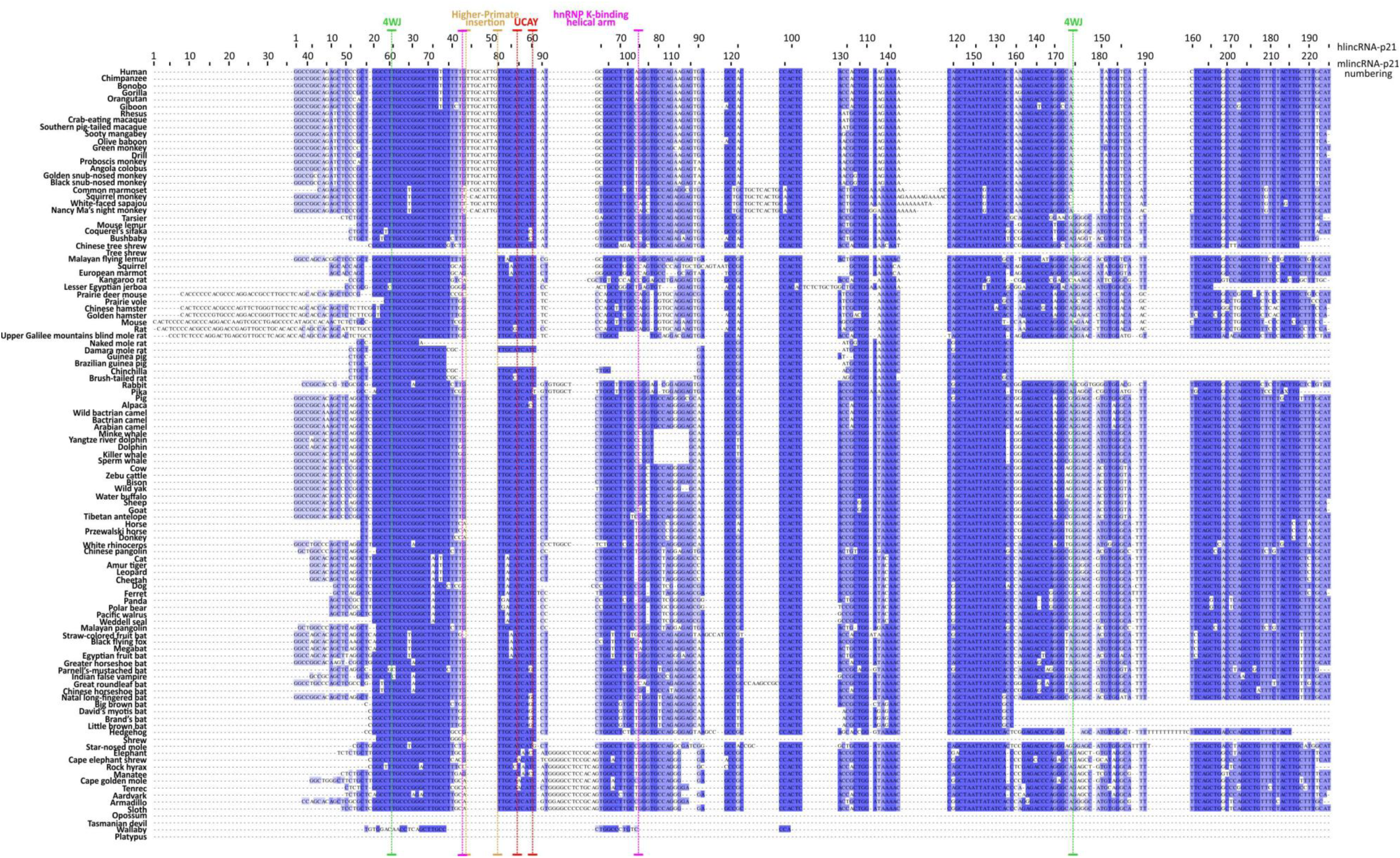
LincRNA-p21 sequence alignment across mammals (related to. Figure 3). Sequence alignment of lincRNA-p21 homologous sequences obtained by BLASTn searches against 111 mammalian species. Sequence alignments were performed by the MUSCLE software through the Jalview cross-platform program. The mammalian species were selected from the USCS Genome Browser PhyloTree 213way.nh (https://github.com/ucscGenomeBrowser/kent/blob/master/src/hg/utils/phyloTrees/213way.nh).

**Figure S5.**
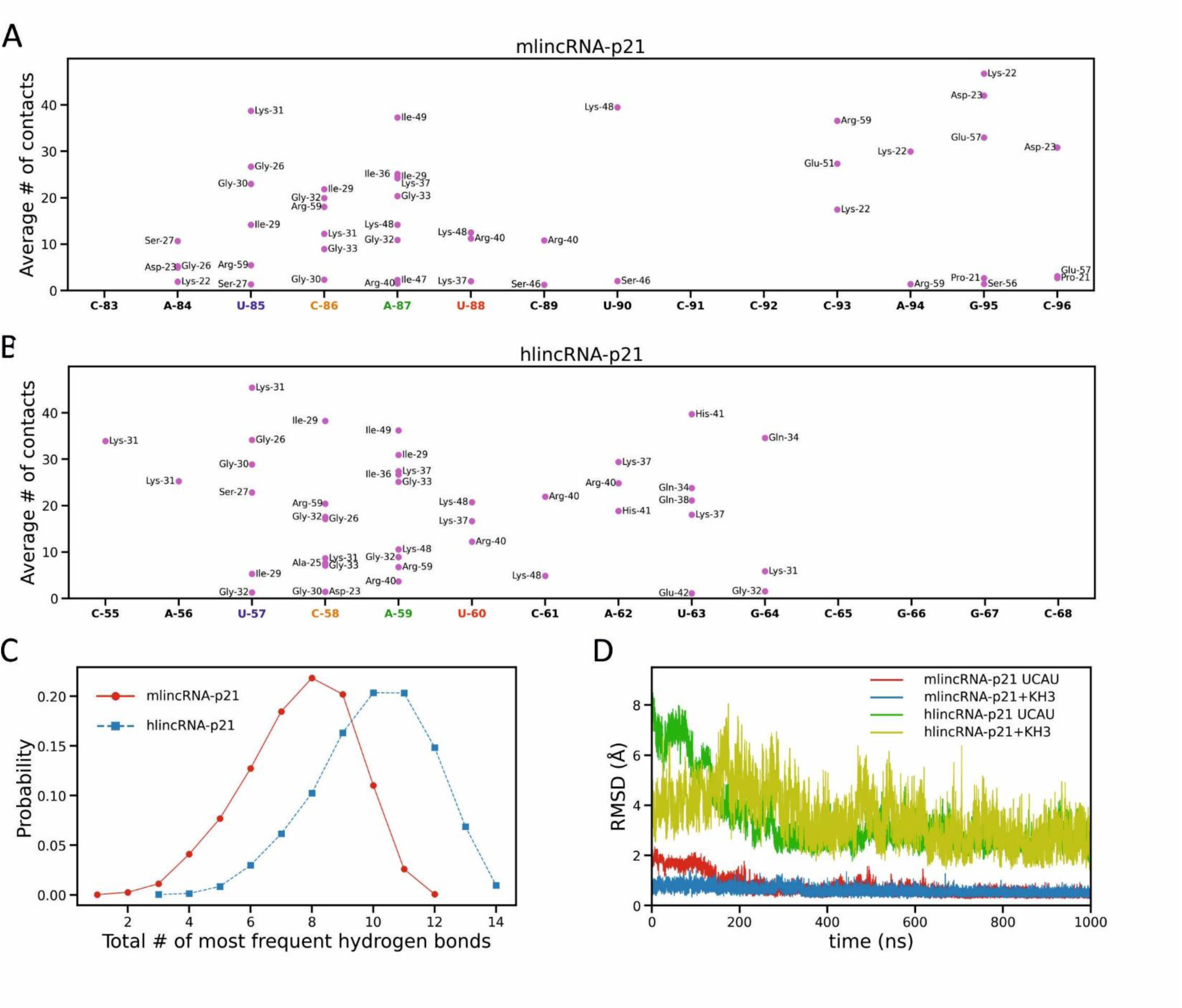
Characterization of the Molecular Dynamics simulations and analysis of the hydrogen bonds and van der Waals interactions established by lincRNA-p21 and the hnRNP K KH3 domain (related to Figure 4). A, B. Maps of van der Waals contacts established by the hnRNP K KH3 domain with mlincRNA-p21 (A) and hlicRNA-p21 (B). The horizontal coordinates correspond to each nucleotide, so that the data are organized in columns with a point for each amino acid in contact with the selected nucleotide. All the points are annotated with the corresponding amino acid. The vertical axes show the average number of atomistic contacts detected for the nucleoside/amino acid couples, based on geometric criteria (distance < 4 Å). The horizontal axes have been shifted to align the tetramer’s location in the two maps. C. Probability distribution of the total number of lincRNA-p21/KH3 hydrogen bonds, where the analysis was limited to the most frequent contacts (>50%), as reported in Fig. 3D and Fig. 3H. Red circles and blue squares correspond to mlincRNA-p21 and hlincRNA-p21, respectively. D. Time dependence of the root-mean-square displacement (RMSD) in simulations of the lincRNA-p21/KH3 complex to assess overall convergence of the simulations. The RMSD is considered for either the UCAU tetramer (red and green lines for hlincRNA-p21 and mlincRNA-p21, respectively) or the full complex (blue and olive for hlincRNA-p21 and mlincRNA-p21, respectively). In all cases, the RMSD is computed using the last frame of the simulation as a reference. The RMSD indicates convergence after about half of the simulation, which supports restricting the analysis to the last 500 ns.

**Figure S6.**
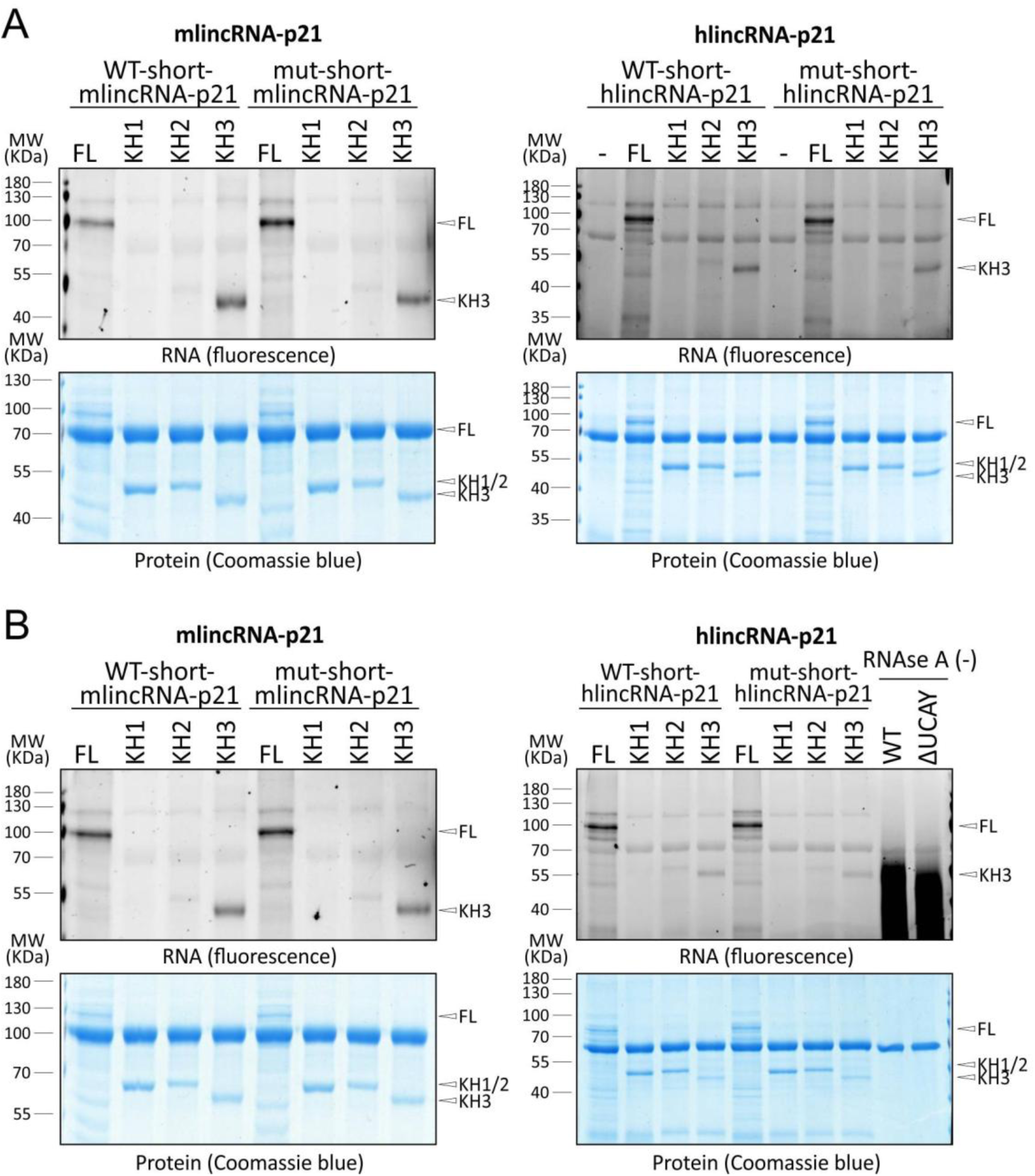
Individual replicates of the protein-RNA UV-crosslinking assay (related to. Figure 5**). A and B.** Individual replicates of the protein-RNA UV-crosslinking assay. Interactions were detected on denaturing gels by Cy5-UTP signal (upper panels). Coomassie-stained gels reveal the protein content (lower panels). BSA is visible at 70 kDa on the Coomassie gel, and a ladder is presented to underscore the corresponding molecular weight. The quantification of the relative RNA-binding capacity of the isolated KH domains compared to the full-length hnRNP K protein is presented in Fig. 5.

**Figure S7.**
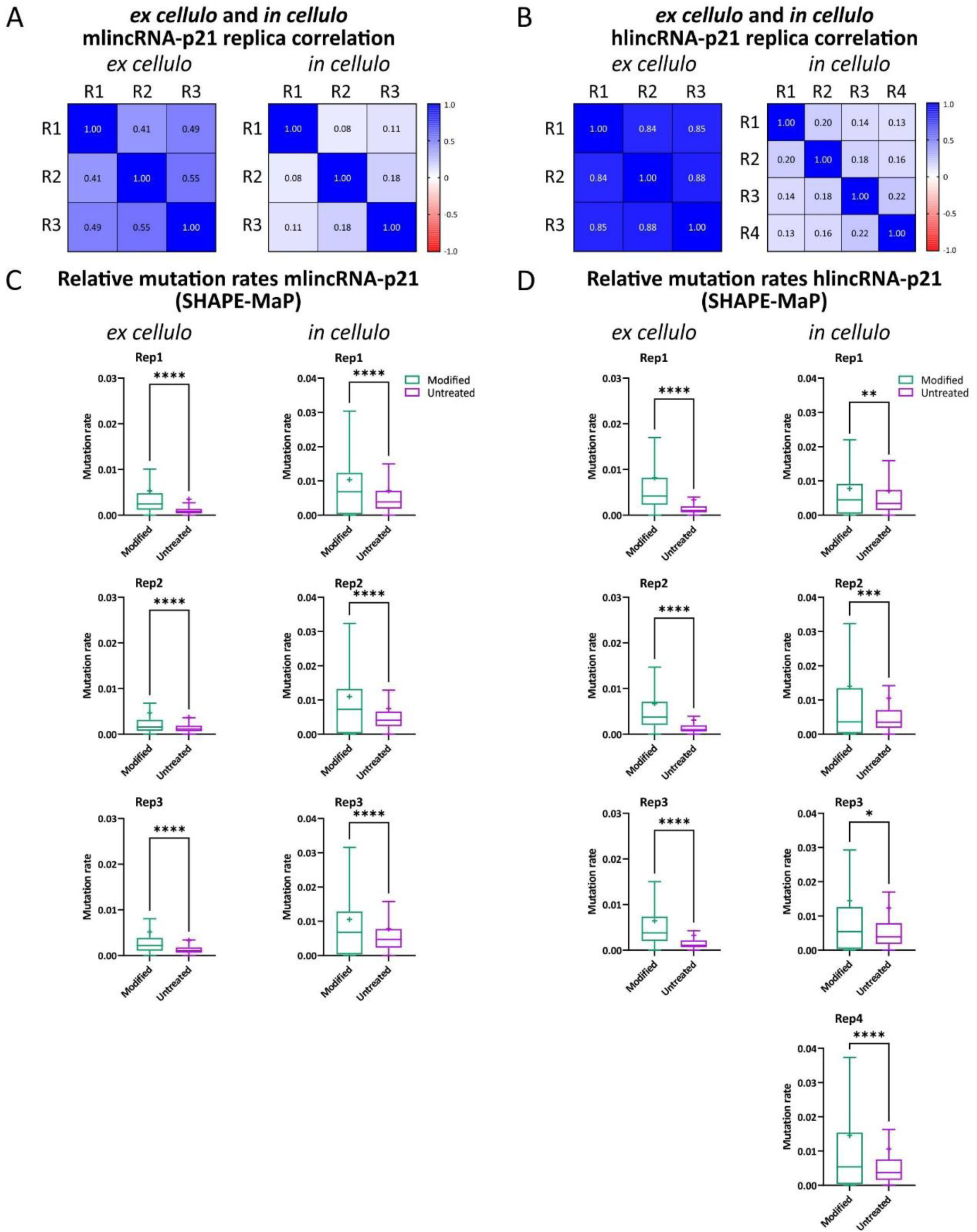
Quality control metrics for the *in cellulo and ex cellulo* SHAPE-MaP (related to. Figure 6**). A and B.** Heat map representation of the *ex cellulo* (left) and *in cellulo* (right) SHAPE-MaP replicate correlation for mlincRNA-p21 (A) and hlincRNA-p21 (B). **C and D.** Mutation rates for the replicates of *in cellulo* (left) and *ex cellulo* (right) mlincRNA-p21 (C) and hlincRNA-p21 (D) (box, interquartile range [IQR]; median indicated by line; average indicated by “x”; whiskers are drawn in the Tukey style, and values outside this range are not shown).

**Figure S8.**
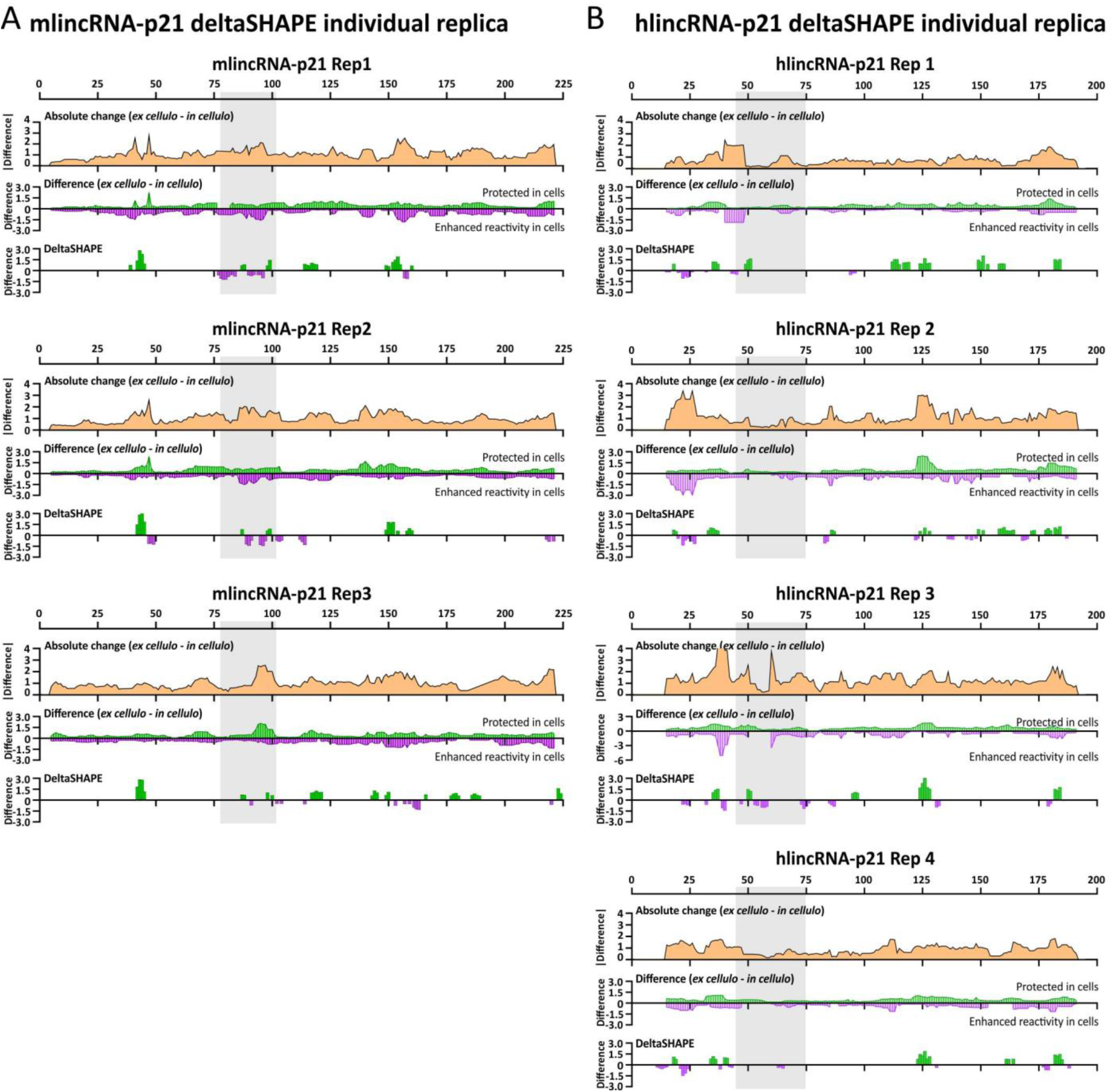
Analysis of the ΔSHAPE individual replicates (related to. Figure 6**). A and B.** ΔSHAPE analyses for each replicate of mlincRNA-p21 (C) and hlincRNA-p21 (D). (Top panel) The absolute change in SHAPE reactivity between *ex cellulo* and *in cellulo* states, smoothed over 50-nt windows. (Middle panel) Contributions of positive and negative differences (red and blue, respectively) to the absolute change, where positive values indicate protection in cells, and negative values indicate enhanced reactivity in cells due to structural reorganization. (Lower panel). ΔSHAPE values indicating regions of the RNA exhibiting statistically significant changes between *in cellulo* and *ex cellulo* conditions. The grey box represents the position of the putative hnRNP K helical arm. The grey box represents the position of the putative hnRNP K helical arm.

**Figure S9.**
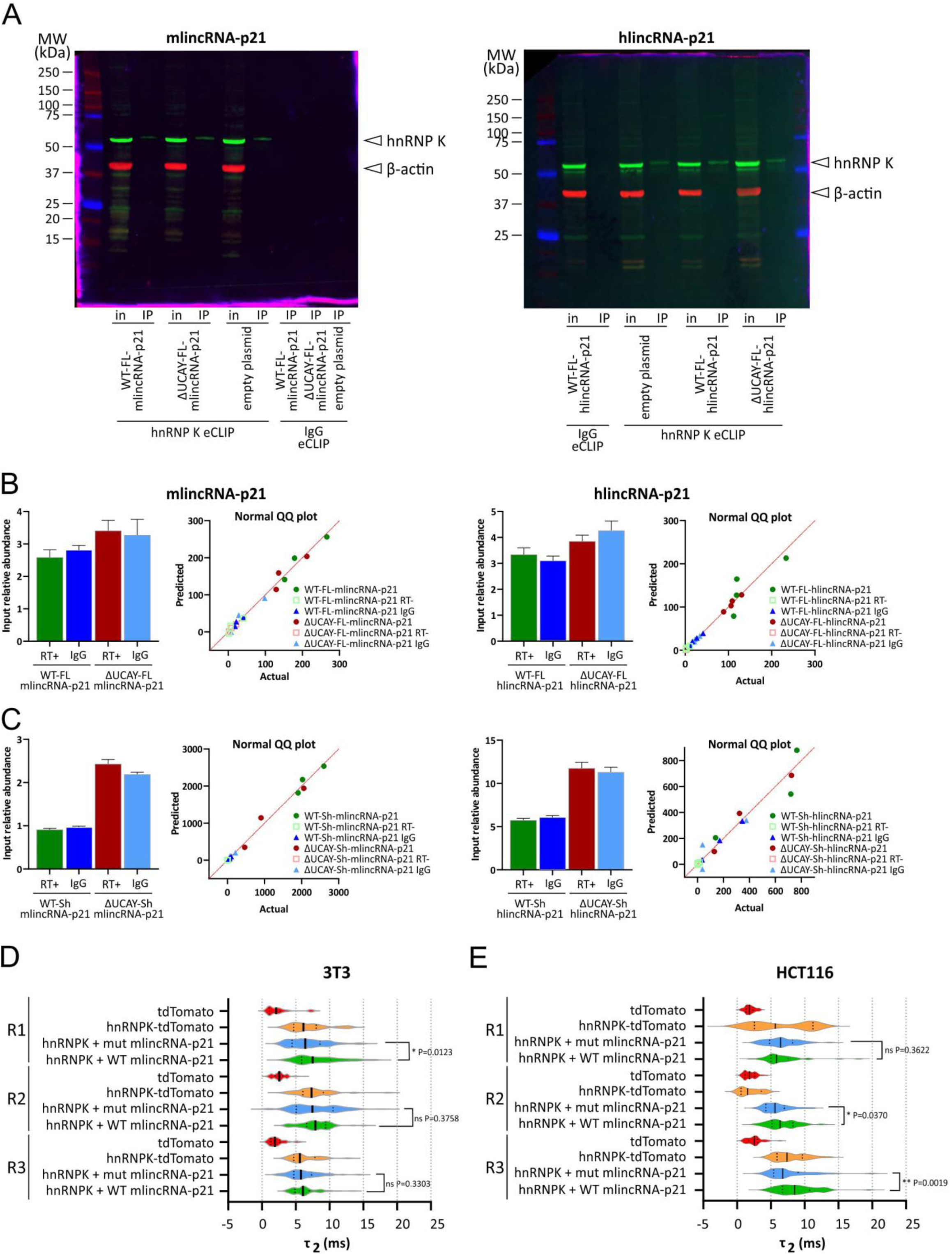
Controls of the eCLIP-qRT-PCR assay and analysis of the FCS individual replicates (Related to. Figure 7**). A.** Western blot analysis of a representative eCLIP-qRT-PCR assay for mlincRNA-p21 (left panel) and hlincRNA-p21 (right panel). The beta-actin protein is only found in the input samples, while the hnRNP K protein is found in both input and IP samples, except for the normal immunoglobulin (IgG) IP samples. **B and C.** (Bar graphs) Quantification by qRT-PCR of the lincRNA-p21 expression in the input samples for each eCLIP-qRT-PCR assay using the FL (B) and short (C) constructs for mlincRNA-p21 (left panels) and hlincRNA-p21 (right panels). (Normal QQ plots) Normality test of both input and IP samples for each eCLIP-qRT-PCR assay using the FL (B) and short (C) constructs for mlincRNA-p21 (left panels) and hlincRNA-p21 (right panels). The QQ plot represents the actual Y values on the horizontal axis, and the predicted Y values (assuming sampling from a Gaussian distribution) on the Y axis. A straight line that matches the line of identity indicates that the data were sampled from a Gaussian (normal) distribution, which is necessary to perform the estimation plot (Fig. 5 B-C). **D and E.** Individual biological replicates of the FCS assays performed in murine 3T3 cells (D) and human HCT116 cells (E). The differences between the WT and mutant constructs as calculated by a Mann-Whitney non-parametric test are displayed for each replicate. Asterisks represent statistically significant differences (P < 0.05).

**Figure S10.**
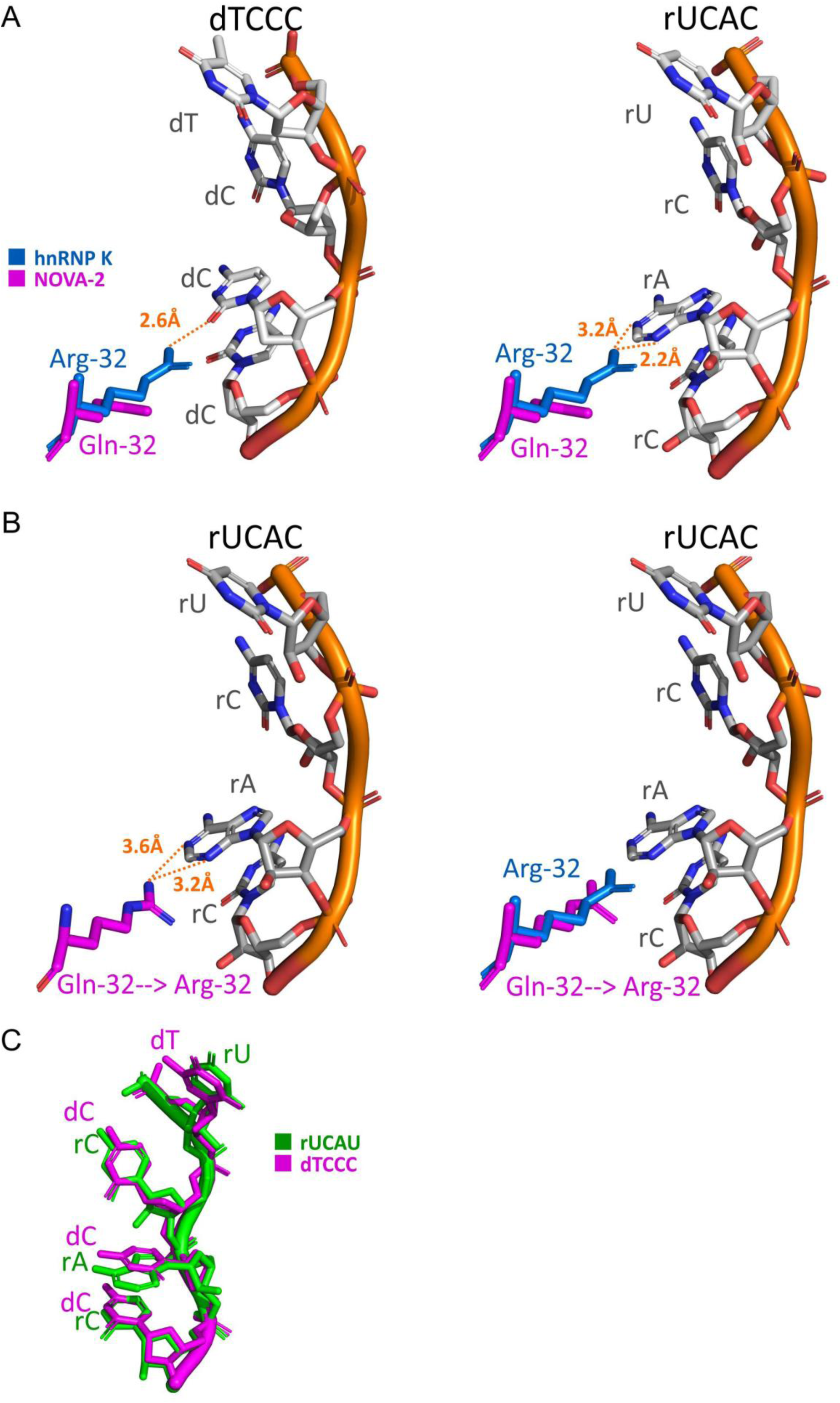
Analysis of the KH3 crystal structure interactions with the third position of the dTCCC and rUCAC ligand tetranucleotides (related to Figure 2). A. Graphical representation of the dTCCC (PDB 1ZZI) (left) and rUCAC (PDB 1EC6) (right) tetranucleotides displaying the distance in Angstroms between the third position of each tetranucleotide and the hnRNP K KH3 arginine-32 (PDB 1ZZI) (blue). B. *In silico* mutation of the NOVA-2 KH3 glutamine-32 (PDB 1EC6) into arginine-32 (present in hnRNP K). (Left panel) Representation of the distance in Angstroms between the mutated arginine-32 (pink) and the adenosine of the rUCAC tetranucleotide (PDB 1EC6). (Right panel) Superposition of the mutated arginine-32 (pink) and the arginine-32 (PDB 1ZZI), showcasing their slightly different spatial localization to the adenosine of the rUCAC tetranucleotide (PDB 1EC6). C. Superposition of the dTCCC and rUCAC ligand tetranucleotides.

**Table S1.**
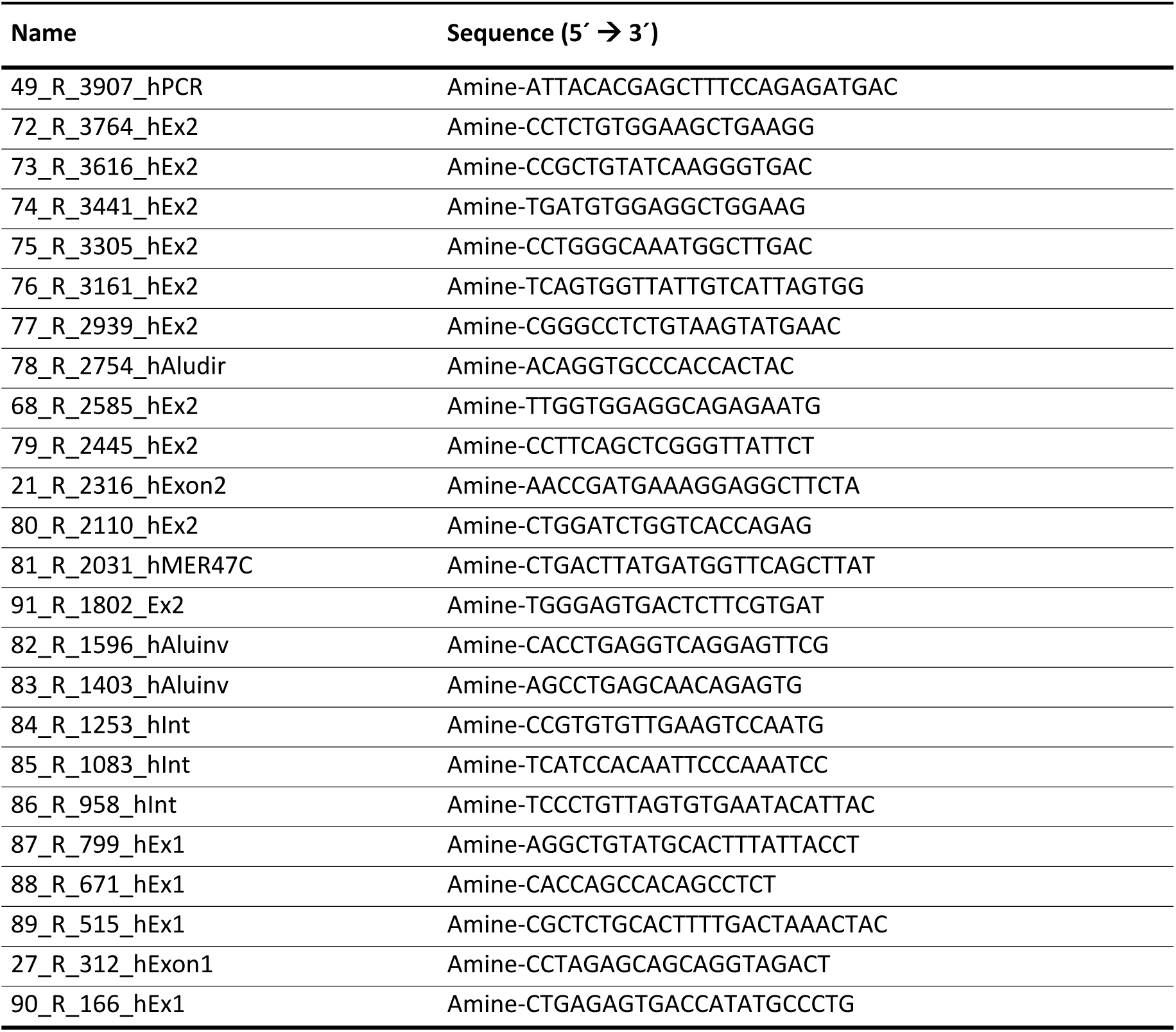
Oligonucleotides used for SHAPE.

**Table S2.**
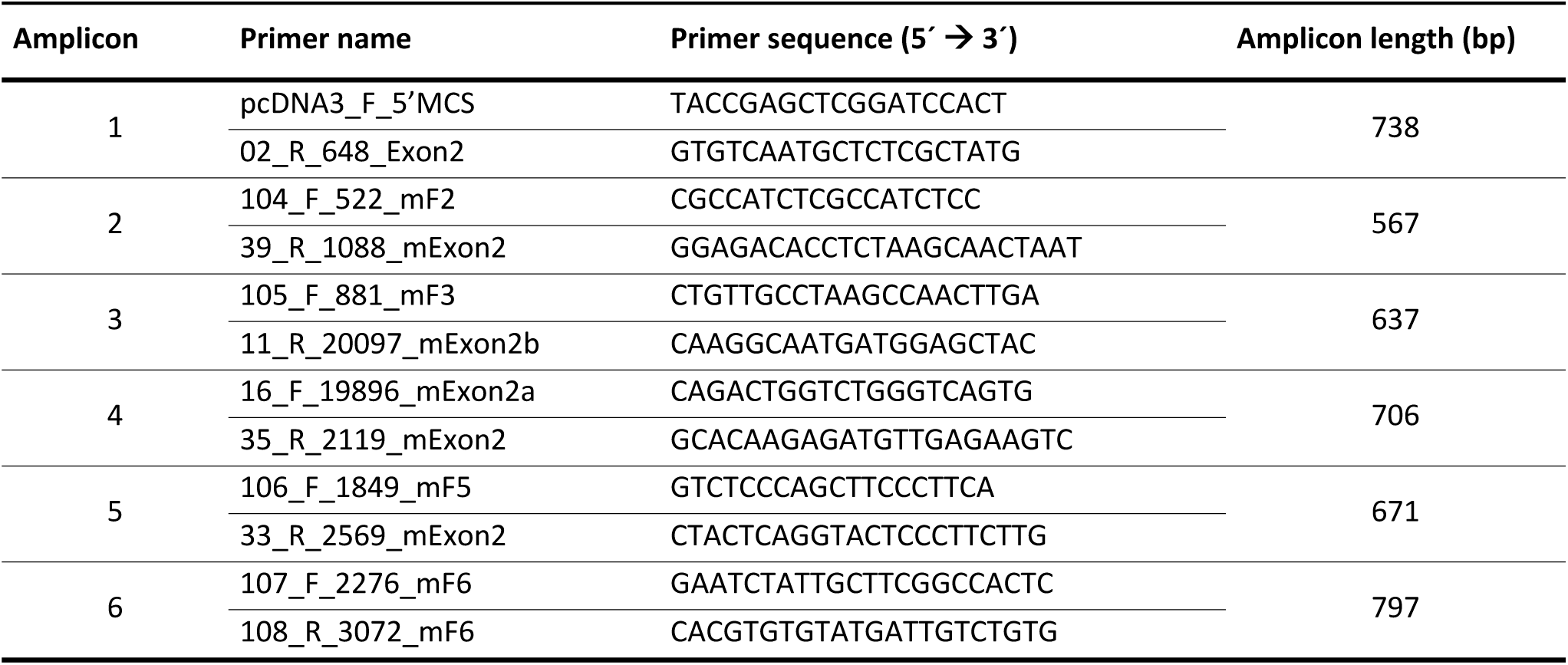
Oligonucleotides used for SHAPE-MaP.

## REFERENCES

1. Unfried, J.P. & Ulitsky, I. Direct roles of long non-coding RNAs in transcription activation. Nat Struct Mol Biol 33, 742–755 (2026).

2. Palazzo, A.F. & Koonin, E.V. Functional Long Non-coding RNAs Evolve from Junk Transcripts. Cell (2020).

3. Camilleri-Robles, C. et al. Genomic and functional conservation of lncRNAs: lessons from flies. Mamm Genome 33, 328–342 (2022).

4. Chillon, I. & Marcia, M. The molecular structure of long non-coding RNAs: emerging patterns and functional implications. Crit Rev Biochem Mol Biol 55, 662–690 (2020).

5. Uroda, T. et al. Conserved Pseudoknots in lncRNA MEG3 Are Essential for Stimulation of the p53 Pathway. Mol Cell 75, 982–995 e9 (2019).

6. Monroy-Eklund, A., Taylor, C., Weidmann, C.A., Burch, C. & Laederach, A. Structural analysis of MALAT1 long noncoding RNA in cells and in evolution. RNA 29, 691–704 (2023).

7. Huarte, M. et al. A large intergenic noncoding RNA induced by p53 mediates global gene repression in the p53 response. Cell 142, 409–19 (2010).

8. Chillon, I. & Pyle, A.M. Inverted repeat Alu elements in the human lincRNA-p21 adopt a conserved secondary structure that regulates RNA function. Nucleic Acids Res 44, 9462–9471 (2016).

9. Dimitrova, N. et al. LincRNA-p21 activates p21 in cis to promote Polycomb target gene expression and to enforce the G1/S checkpoint. Mol Cell 54, 777–90 (2014).

10. Winkler, L. et al. Functional elements of the cis-regulatory lincRNA-p21. Cell Rep 39, 110687 (2022).

11. Wu, G. et al. LincRNA-p21 regulates neointima formation, vascular smooth muscle cell proliferation, apoptosis, and atherosclerosis by enhancing p53 activity. Circulation 130, 1452–1465 (2014).

12. Weeks, K.M. SHAPE Directed Discovery of New Functions in Large RNAs. Acc Chem Res 54, 2502–2517 (2021).

13. Chillon, I. et al. Native Purification and Analysis of Long RNAs. Methods Enzymol 558, 3–37 (2015).

14. Somarowthu, S. et al. HOTAIR forms an intricate and modular secondary structure. Mol Cell 58, 353–61 (2015).

15. Rice, G.M., Leonard, C.W. & Weeks, K.M. RNA secondary structure modeling at consistent high accuracy using differential SHAPE. RNA 20, 846–54 (2014).

16. Smola, M.J., Rice, G.M., Busan, S., Siegfried, N.A. & Weeks, K.M. Selective 2’-hydroxyl acylation analyzed by primer extension and mutational profiling (SHAPE-MaP) for direct, versatile and accurate RNA structure analysis. Nat Protoc 10, 1643–69 (2015).

17. Dethoff, E.A. & Weeks, K.M. Effects of Refolding on Large-Scale RNA Structure. Biochemistry 58, 3069-3077 (2019).

18. Smola, M.J. et al. SHAPE reveals transcript-wide interactions, complex structural domains, and protein interactions across the Xist lncRNA in living cells. Proc Natl Acad Sci U S A 113, 10322–7 (2016).

19. Mauger, D.M. et al. Functionally conserved architecture of hepatitis C virus RNA genomes. Proc Natl Acad Sci U S A 112, 3692–7 (2015).

20. Watts, J.M. et al. Architecture and secondary structure of an entire HIV-1 RNA genome. Nature 460, 711–6 (2009).

21. Reuter, J.S. & Mathews, D.H. RNAstructure: software for RNA secondary structure prediction and analysis. BMC Bioinformatics 11, 129 (2010).

22. Jensen, K.B., Musunuru, K., Lewis, H.A., Burley, S.K. & Darnell, R.B. The tetranucleotide UCAY directs the specific recognition of RNA by the Nova K-homology 3 domain. Proc Natl Acad Sci U S A 97, 5740–5 (2000).

23. Buckanovich, R.J. & Darnell, R.B. The neuronal RNA binding protein Nova-1 recognizes specific RNA targets in vitro and in vivo. Mol Cell Biol 17, 3194–201 (1997).

24. Valverde, R., Edwards, L. & Regan, L. Structure and function of KH domains. FEBS J 275, 2712–26 (2008).

25. Lewis, H.A. et al. Sequence-specific RNA binding by a Nova KH domain: implications for paraneoplastic disease and the fragile X syndrome. Cell 100, 323–32 (2000).

26. Backe, P.H., Messias, A.C., Ravelli, R.B., Sattler, M. & Cusack, S. X-ray crystallographic and NMR studies of the third KH domain of hnRNP K in complex with single-stranded nucleic acids. Structure 13, 1055–67 (2005).

27. Boniecki, M.J. et al. SimRNA: a coarse-grained method for RNA folding simulations and 3D structure prediction. Nucleic Acids Res 44, e63 (2016).

28. Moafinejad, S.N. et al. SimRNAweb v2.0: a web server for RNA folding simulations and 3D structure modeling, with optional restraints and enhanced analysis of folding trajectories. Nucleic Acids Res 52, W368–W373 (2024).

29. Case, D.A. et al. AmberTools. Journal of Chemical Information and Modeling 63, 6183–6191 (2023).

30. Smola, M.J., Calabrese, J.M. & Weeks, K.M. Detection of RNA-Protein Interactions in Living Cells with SHAPE. Biochemistry 54, 6867–75 (2015).

31. Wang, Z. et al. The emerging roles of hnRNPK. J Cell Physiol 235, 1995–2008 (2020).

32. Herrera-Ubeda, C. et al. Microsyntenic Clusters Reveal Conservation of lncRNAs in Chordates Despite Absence of Sequence Conservation. Biology (Basel) 8(2019).

33. Olazagoitia-Garmendia, A., Senovilla-Ganzo, R., García-Moreno, F. & Castellanos-Rubio, A. Functional evolutionary convergence of long noncoding RNAs involved in embryonic development. Communications Biology 6, 908 (2023).

34. Seemann, S.E. et al. The identification and functional annotation of RNA structures conserved in vertebrates. Genome Res 27, 1371–1383 (2017).

35. Corley, M., Burns, M.C. & Yeo, G.W. How RNA-Binding Proteins Interact with RNA: Molecules and Mechanisms. Mol Cell 78, 9–29 (2020).

36. Cech, T.R., Davidovich, C. & Jenner, R.G. PRC2-RNA interactions: Viewpoint from Tom Cech, Chen Davidovich, and Richard Jenner. Mol Cell 84, 3593–3595 (2024).

37. Yoon, J.H. et al. LincRNA-p21 suppresses target mRNA translation. Mol Cell 47, 648–55 (2012).

38. Yang, F., Zhang, H., Mei, Y. & Wu, M. Reciprocal regulation of HIF-1alpha and lincRNA-p21 modulates the Warburg effect. Mol Cell 53, 88–100 (2014).

39. Trotman, J.B. et al. Isogenic comparison of Airn and Xist reveals core principles of Polycomb recruitment by lncRNAs. Mol Cell 85, 1117–1133 e14 (2025).

40. Nakamoto, M.Y., Lammer, N.C., Batey, R.T. & Wuttke, D.S. hnRNPK recognition of the B motif of Xist and other biological RNAs. Nucleic Acids Research 48, 9320–9335 (2020).

41. Ostrowski, J., Wyrwicz, L., Rychlewski, L. & Bomsztyk, K. Heterogeneous Nuclear Ribonucleoprotein K Protein Associates with Multiple Mitochondrial Transcripts within the Organelle *. Journal of Biological Chemistry 277, 6303–6310 (2002).

42. Dominguez, D., et al. Sequence, Structure, and Context Preferences of Human RNA Binding Proteins. Mol Cell 70, 854–867 e9 (2018).

43. Thisted, T., Lyakhov, D.L. & Liebhaber, S.A. Optimized RNA targets of two closely related triple KH domain proteins, heterogeneous nuclear ribonucleoprotein K and alphaCP-2KL, suggest Distinct modes of RNA recognition. J Biol Chem 276, 17484–96 (2001).

44. Kribelbauer, J.F., Rastogi, C., Bussemaker, H.J. & Mann, R.S. Low-Affinity Binding Sites and the Transcription Factor Specificity Paradox in Eukaryotes. Annu Rev Cell Dev Biol 35, 357–379 (2019).

45. Pintacuda, G. et al. hnRNPK Recruits PCGF3/5-PRC1 to the Xist RNA B-Repeat to Establish Polycomb-Mediated Chromosomal Silencing. Mol Cell 68, 955–969 e10 (2017).

46. Karabiber, F., McGinnis, J.L., Favorov, O.V. & Weeks, K.M. QuShape: rapid, accurate, and best-practices quantification of nucleic acid probing information, resolved by capillary electrophoresis. RNA 19, 63–73 (2013).

47. Siegfried, N.A., Busan, S., Rice, G.M., Nelson, J.A. & Weeks, K.M. RNA motif discovery by SHAPE and mutational profiling (SHAPE-MaP). Nat Methods 11, 959–65 (2014).

48. Mustoe, A.M., Lama, N.N., Irving, P.S., Olson, S.W. & Weeks, K.M. RNA base-pairing complexity in living cells visualized by correlated chemical probing. Proc Natl Acad Sci U S A 116, 24574–24582 (2019).

49. Busan, S. & Weeks, K.M. Accurate detection of chemical modifications in RNA by mutational profiling (MaP) with ShapeMapper 2. RNA 24, 143–148 (2018).

50. Darty, K., Denise, A. & Ponty, Y. VARNA: Interactive drawing and editing of the RNA secondary structure. Bioinformatics 25, 1974–5 (2009).

51. Smola, M.J. & Weeks, K.M. In-cell RNA structure probing with SHAPE-MaP. Nature Protocols 13, 1181–1195 (2018).

52. Wang, J., Cieplak, P. & Kollman, P.A. How well does a restrained electrostatic potential (RESP) model perform in calculating conformational energies of organic and biological molecules? Journal of Computational Chemistry 21, 1049–1074 (2000).

53. Perez, A. et al. Refinement of the AMBER force field for nucleic acids: improving the description of alpha/gamma conformers. Biophys J 92, 3817–29 (2007).

54. Zgarbova, M. et al. Refinement of the Cornell et al. Nucleic Acids Force Field Based on Reference Quantum Chemical Calculations of Glycosidic Torsion Profiles. J Chem Theory Comput 7, 2886–2902 (2011).

55. Jorgensen, W.L., Chandrasekhar, J., Madura, J.D., Impey, R.W. & Klein, M.L. Comparison of simple potential functions for simulating liquid water. The Journal of Chemical Physics 79, 926–935 (1983).

56. Joung, I.S. & Cheatham, T.E., 3rd. Molecular dynamics simulations of the dynamic and energetic properties of alkali and halide ions using water-model-specific ion parameters. J Phys Chem B 113, 13279–90 (2009).

57. Maier, J.A. et al. ff14SB: Improving the Accuracy of Protein Side Chain and Backbone Parameters from ff99SB. J Chem Theory Comput 11, 3696–713 (2015).

58. Kaus, J.W., Pierce, L.T., Walker, R.C. & McCammont, J.A. Improving the Efficiency of Free Energy Calculations in the Amber Molecular Dynamics Package. J Chem Theory Comput 9(2013).

59. Roe, D.R. & Cheatham, T.E., 3rd. PTRAJ and CPPTRAJ: Software for Processing and Analysis of Molecular Dynamics Trajectory Data. J Chem Theory Comput 9, 3084–95 (2013).

60. Frenkel, D. & Smit, B. Understanding molecular simulation : from algorithms to applications, xviii, 443 p. (Academic Press, San Diego, 1996).

61. Madeira, F. et al. The EMBL-EBI Job Dispatcher sequence analysis tools framework in 2024. Nucleic Acids Res> 52, W521-W525 (2024).

62. Waterhouse, A.M., Procter, J.B., Martin, D.M., Clamp, M. & Barton, G.J. Jalview Version 2--a multiple sequence alignment editor and analysis workbench. Bioinformatics 25, 1189–91 (2009).

63. Nawrocki, E.P., Kolbe, D.L. & Eddy, S.R. Infernal 1.0: inference of RNA alignments. Bioinformatics 25, 1335–7 (2009).

64. Rivas, E., Clements, J. & Eddy, S.R. Estimating the power of sequence covariation for detecting conserved RNA structure. Bioinformatics (2020).

65. Rivas, E. RNA covariation at helix-level resolution for the identification of evolutionarily conserved RNA structure. PLoS Comput Biol 19, e1011262 (2023).

66. Carnesecchi, J. et al. The Hox transcription factor Ultrabithorax binds RNA and regulates co-transcriptional splicing through an interplay with RNA polymerase II. Nucleic Acids Res 50, 763–783 (2022).

67. Van Nostrand, E.L. et al. Robust transcriptome-wide discovery of RNA-binding protein binding sites with enhanced CLIP (eCLIP). Nat Methods 13, 508–14 (2016).

